# Conserved Methyltransferase Spb1 Targets mRNAs for Regulated Modification with 2′-O-Methyl Ribose

**DOI:** 10.1101/271916

**Authors:** Kristen M. Bartoli, Cassandra Schaening, Thomas M. Carlile, Wendy V. Gilbert

## Abstract

Non-coding RNAs contain dozens of chemically distinct modifications, of which only a few have been identified in mRNAs. The recent discovery that certain tRNA modifying enzymes also target mRNAs suggests the potential for many additional mRNA modifications. Here, we show that conserved tRNA 2′-O-methyltransferases Trm3, 7,13 and 44, and rRNA 2′-O-methyltransferase Spb1, interact with specific mRNA sites in yeast by crosslinking immunoprecipitation and sequencing (CLIP-seq). We developed sequencing of methylation at two prime hydroxyls (MeTH-seq) for transcriptome-wide mapping of 2′-O-methyl ribose (Nm) with single-nucleotide resolution, and discover thousands of potential Nm sites in mRNAs. Genetic analysis identified hundreds of mRNA targets for the Spb1 methyltransferase, which can target both mRNA and non-coding RNA for environmentally regulated modification. Our work identifies Nm as a prevalent mRNA modification that is likely to be conserved and provides methods to investigate its distribution and regulation.

**HIGHLIGHTS:**

- MeTH-seq identifies 2′-O-methylribose genome-wide at single-nucleotide resolution
- Five conserved methyltransferases interact with yeast mRNA
- Spb1 is a major mRNA 2′-O-methyltransferase, and targets most ribosomal protein mRNAs
- *SPB1* expression is required to maintain normal levels of Spb1 target mRNAs

## INTRODUCTION

Non-coding RNAs, in particular transfer RNAs (tRNA), are heavily modified with more than 100 distinct post-transcriptional modifications that affect RNA structure, stability, and function (Machnicka et al., 2013). The recent development of protocols for transcriptome-wide mapping of certain RNA modifications has revealed a complex and dynamic mRNA ‘epitranscriptome’ that includes N6-methyladenosine (m^6^A), 5-methylcytidine (m^5^C), inosine (I), pseudouridine (Ψ), 5-hydroxymethylcytidine (hm^5^C), and N1-methyladenosine (m^1^A) in eukaryotic cells (Gilbert et al., 2016; Li et al., 2016; Schaefer et al., 2017). Emerging evidence links regulated mRNA modifications, in particular m^6^A, to post-transcriptional gene regulation in diverse biological systems (Roignant and Soller, 2017; Yue et al., 2015). However, the full extent and diversity of mRNA modifications is unknown.

The discovery that some tRNA modifying enzymes, specifically pseudouridine synthases (Pus), also target mRNAs (Carlile et al., 2014; Lovejoy et al., 2014; Schwartz et al., 2014), suggests mRNAs might contain many more modifications than are currently known. Notably, the conserved tRNA 2′-O-methyltransferase, Trm44, was identified by proteomic analysis of highly purified UV cross-linked poly(A)^+^ RNPs in yeast (Beckmann et al., 2015), together with Pus1 and Pus7, which modify dozens of mRNA targets (Carlile et al., 2014; Schwartz et al., 2014). Although no specific mRNA binding sites were identified for Trm44, its association with polyadenylated RNA in vivo suggests it may also target mRNAs in addition to tRNAs.

2′-O-methyl ribose can occur on any base (Nm) and is an abundant and highly conserved modification found at multiple locations in tRNA (Figure S1A), ribosomal RNA (rRNA), and small nuclear RNA (snRNA). Yeast tRNAs are 2′-O-methylated by four conserved Trm proteins. Trm3, orthologous to human TARBP1, modifies G18 or G19 in the D-loop of 12 different tRNAs (Cavaillé et al., 1999). Trm7 (FTSJ1) modifies two positions in the anticodon loops of tRNA^Phe^, tRNA^Trp^, and tRNA^Leu^ (Pintard et al., 2002a). Trm13 (CCDC76) modifies tRNA^Gly^, tRNA^His^, and tRNA^Pro^ in the acceptor stem at position 4 or 5 (Wilkinson et al., 2007), and Trm44 (METTL19) modifies tRNA^Ser^ at position 44 or 45 in the variable loop (Kotelawala et al., 2008). Four additional conserved 2′-O-methyltranserases target cytosolic and mitochondrial ribosomes. Three of these are protein-only enzymes: Spb1 (FTSJ3), which modifies Gm2922 of the 25S rRNA (Lapeyre and Purushothaman, 2004), and Mrm1 (MRM1) and Mrm2 (MRM2), which modify Gm2270 and Um2791, respectively, of the 21S rRNA (Pintard et al., 2002b; Sirum-Connolly and Mason, 1993). The fourth, Nop1 (FBL), forms RNP complexes containing additional proteins and C/D-box small nucleolar RNAs (snoRNA) that direct methylation of dozens of sites on the 18S and 25S rRNAs by base pairing with the guide RNA (Kiss-László et al., 1996; Kiss, 2002). Unlike the snoRNA-guided enzymes, the basis for substrate specificity is incompletely understood for the protein-only 2′-O-methyltransferases. Thus, it is not possible to predict their RNA targets by bioinformatic analysis.

2′-O-methylation – and regulation of this modification – has the potential to broadly affect mRNA metabolism through effects on RNA structure, stability, RNA-protein interactions, and translation. 2′-O-methyl ribose increases the thermodynamic stability of RNA:RNA base pairs and stabilizes the C3′ endo conformation of ribose found in A-form RNA duplexes (Inoue et al., 1987; Kawai et al., 1992; Majlessi et al., 1998; Tsourkas et al., 2002). In addition, many RNA tertiary structures involve the 2′ hydroxyl groups of ribose (Butcher and Pyle, 2011), which can be disrupted by 2′-O-methylation (Lebars et al., 2008). Methylation of 2′ hydroxyls can also inhibit RNA-protein interactions through steric effects (Hou et al., 2001; Lacoux et al., 2012) or by impacting hydrogen bonding, which frequently involves the 2′ hydroxyl group (Jones et al., 2001; Treger and Westhof, 2001). 2′-O-methylated nucleotides are also resistant to a variety of nucleases (Sproat et al., 1989). Finally, 2′-O-methylation of synthetic mRNAs interferes with translation and induces site-specific ribosome stalling in vitro (Dunlap et al., 1971; Hoernes et al., 2015). Thus, it is of particular interest to know which natural mRNAs contain Nm and what enzymes install this modification in messenger RNAs.

Recent evidence suggests 2′-O-methyl ribose could be abundant in human mRNA, but the methyltransferases were not identified (Dai et al., 2017). Here we determine mRNA interaction sites genome-wide for five conserved RNA 2′-O-methyltransferases in yeast by UV crosslinking and immunoprecipitation (CLIP-seq). We then develop MeTH-seq, a genome-wide, single-nucleotide-resolution method to determine the locations of 2′-O-methyl ribose (Nm). We validate MeTH-seq by detecting known sites in non-coding RNAs and conservatively identify 690 novel methyltransferase-dependent Nm sites in yeast mRNAs by comprehensive profiling following deletion or depletion of conserved methyltransferases Trm3, 7, 13, and 44, which canonically target tRNA, and Spb1, which targets rRNA. We identify novel targets for each methyltransferase and find that Spb1 is the predominant mRNA methyltransferase in yeast. We further show that *SPB1* is required to maintain normal expression levels of a coherent regulon encoding proteins necessary for ribosome assembly. Finally, we show that mRNA Nm sites are regulated in response to environmental signals. Our results identify Spb1 as the primary methyltransferase installing Nm as a widespread, regulated mRNA modification that is likely to be conserved from yeast to man.

## RESULTS

### Trm44 tRNA Methyltransferase Interacts with Specific mRNAs In Vivo

The yeast genome encodes more than 60 proteins characterized as tRNA modifying enzymes (Phizicky and Hopper, 2010), of which several have recently been shown to target mRNAs (Carlile et al., 2014; Lovejoy et al., 2014; Schwartz et al., 2014). We hypothesized that mRNAs may be recognized and modified by additional enzymes among this cohort. As a first step towards identifying likely mRNA modifying factors, we examined published mRNA interactome data obtained by UV crosslinking of living yeast cells followed by stringent purification of poly(A)^+^ mRNPs and mass spectrometry (Beckmann et al., 2015). Just four tRNA modifying enzymes – Pus1, Pus7, Dus3 and Trm44 – were found to be mRNA-associated. Two of these, the pseudouridine synthases Pus1 and Pus7, are known to modify dozens of mRNA targets (Carlile et al., 2014; Schwartz et al., 2014). In contrast, the known targets of Dus3 and Trm44 are restricted to tRNAs. Given the broad potential for mRNA 2′-O-methylation to affect mRNA metabolism, we selected Trm44 for further study.

To identify specific mRNAs that interact with Trm44, we performed in vivo UV crosslinking (CL) followed by limited RNase digestion, immunoprecipitation (IP) of HPM-tagged protein expressed from the endogenous *TRM44* locus (Trm44-HPM; Table S1), and Illumina sequencing of the associated RNA fragments (Figure 1A; Methods). As expected, radiolabeling of the crosslinked immunoprecipitated RNA-protein complexes (RNP) showed a prominent band of the predicted size for Trm44-HPM bound to an intact tRNA, which was not present in control IPs from an untagged strain (Figure S1B). We generated CLIP libraries from these samples using an enhanced protocol (eCLIP-Seq) that increases the number of non-PCR duplicate reads (Van Nostrand et al., 2016). In parallel, we prepared a size-matched input (SMI) library to control for abundant RNA fragments that contribute to non-specific background signal (Methods). This comparison is important for identifying true binding sites as normalization of eCLIP reads to SMI has been shown to eliminate >90% of putative RBP binding sites as false positives (Conway et al., 2016; Van Nostrand et al., 2016).

**Figure 1.**
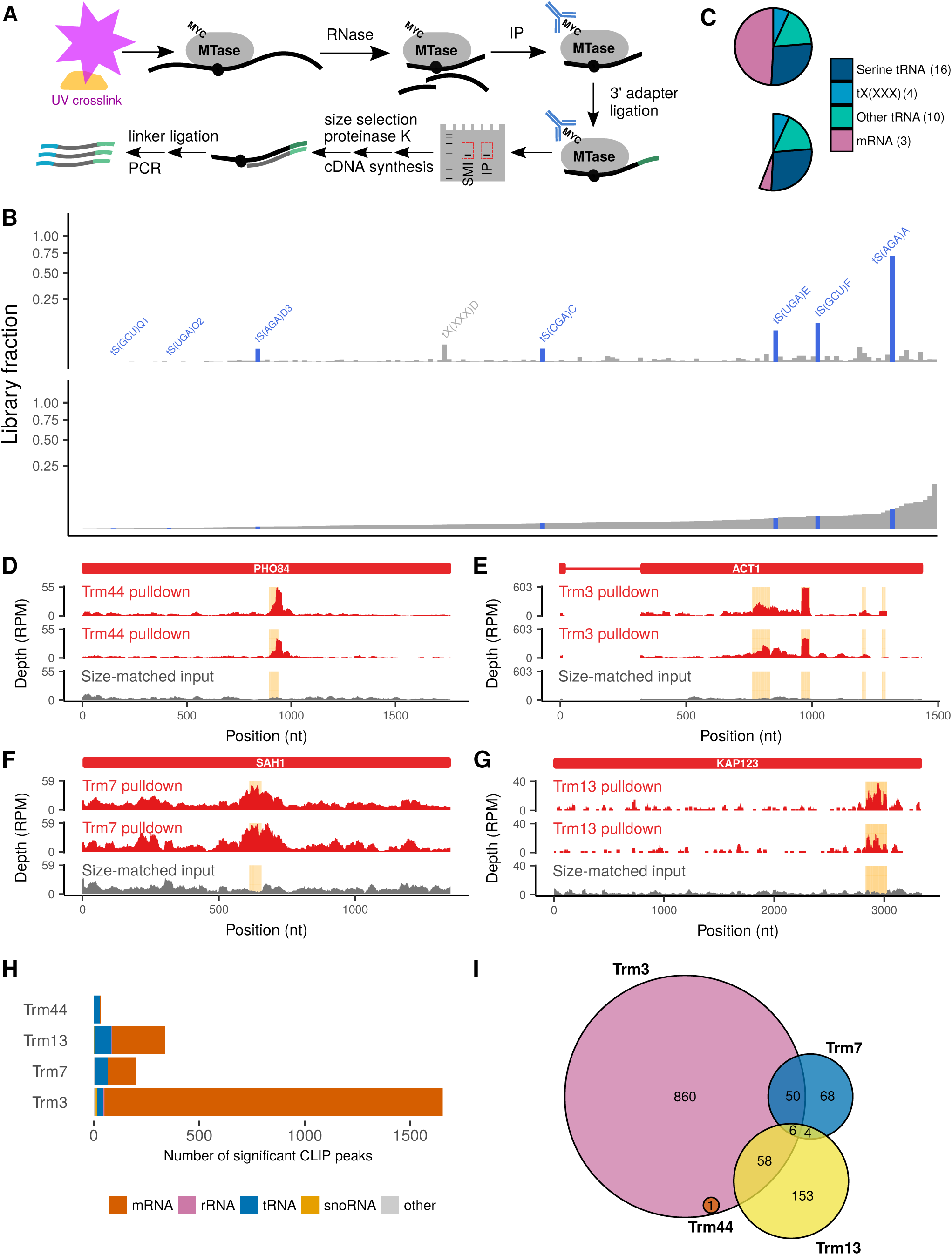
tRNA 2′-O-methyltransferases Associate with mRNAs In Vivo. (A)Schematic of eCLIP procedure on an HPM-tagged methyltransferase (MTase) (B)Overlap between Trm44 non-coding RNA eCLIP targets and methylation substrates. The fraction of the library mapped to each feature is shown for two eCLIP replicates and SMI control. Genes are ordered by their abundance in SMI. tRNA^Ser^ that are known to be modified by Trm44 are highlighted in blue. tRNA^XXX^ (*) is a computationally predicted tRNA^Ser^ of unknown modification status (Lowe and Eddy, 1997). (C)Distribution of significantly enriched eCLIP peaks for Trm44. Top: ≥ 4-fold versus SMI. Bottom: *p* ≤ 0.001 and ≥ 4-fold-enriched versus SMI. Numbers for the bottom pie chart indicated in parentheses. (D-G) Examples of mRNA eCLIP peaks for Trm44 (D), Trm3 (E), Trm7 (F), and Trm13 (G). (H)Number of significant eCLIP peaks in tRNA, mRNA, rRNA, and snoRNA for Trm3, 7, 13 and 44; *p* ≤ 0.001 and ≥4-fold enriched versus SMI. (I)Venn diagram illustrates lack of overlap between mRNAs CLIPed to different Trm proteins.

Consistent with the known methylation targets of Trm44, reads mapping to tRNA were enriched >10-fold in the Trm44-HPM eCLIP libraries compared to SMI while reads mapping to rRNA were depleted >50-fold (Table S2). Among tRNA eCLIP reads there was a striking enrichment for 6 tRNAs compared to SMI: 5 tRNA^Ser^ that are known to be methylated by Trm44 and 1 computationally predicted tRNA of undetermined specificity that is very similar to serine tRNAs (Figure 1B) (Kotelawala et al., 2007; Lowe and Eddy, 1997). Next we applied the CLIPper algorithm (Lovci et al., 2013) to identify clusters of CLIP-Seq reads and computed the enrichment of each cluster in IP compared to SMI (Methods). Using stringent enrichment criteria (p ≤ 0.001 and ≥4-fold enriched versus SMI), we identified 33 high-confidence Trm44 binding sites, 60% of which were located in various tRNA^Ser^. 3 stringent clusters were identified in mRNA, with 26 additional mRNA clusters showing enrichment at relaxed cutoffs (Figure 1C; Table S3). As a further control, we compared Trm44-HPM eCLIP peaks to Pus1-HPM, which bound a large set of tRNAs as expected and did not significantly enrich the Trm44 mRNA binding sites (Figure S1C; Table S4). These results demonstrate that Trm44 interacts with specific mRNA sites in vivo and suggests that mRNA could be 2′-O-methylated.

### Discovery of mRNA Binding Sites for Trm3, Trm7 and Trm13 by eCLIP-Seq

Because mRNP interactome analysis did not identify all tRNA modifying pseudouridine synthases with known mRNA pseudouridylation targets, we reasoned that additional tRNA 2′-O-methyltransferases might associate with specific mRNAs in vivo. We therefore performed eCLIP-Seq on yeast strains expressing HPM-tagged versions of Trm3, Trm7 and Trm13 under their native promoters with SMI controls for each (Figure S1D). As expected, tRNA mapping reads were enriched in these eCLIP libraries (Table S2). CLIPper peak discovery and normalization to SMI identified stringent peaks for each Trm protein (p ≤ 0.001 and ≥4-fold enriched versus SMI; Methods), and most known methylation targets were enriched including tRNA^His^, tRNA^Leu^, tRNA^Ser^, and tRNA^Tyr^ for Trm3; tRNA^Leu^, tRNA^Phe^, and tRNA^Trp^ for Trm7; and tRNA^Gly^, tRNA^His^, and tRNA^Pro^ for Trm13 (Figures S1E-S1G; Tables S5-S7). In addition, these eCLIPs each produced numerous stringent peaks that mapped to specific mRNA sites distinct from one another and from Trm44-HPM (Figures 1D-1I; Tables S3 and S5-S7). Thus, each of the tRNA 2′-O-methyltransferases interacts with specific mRNAs in vivo in addition to their canonical tRNA targets.

We also identified stringent peaks in non-coding RNAs not known to be methylation targets of the Trm enzymes. A substantial fraction of significantly enriched non-coding RNA peaks occurred in other tRNAs (Tables S3 and S5-S7). Moreover, in a few cases, such as tRNA^Ser^ crosslinking to Trm7 and tRNA^Val^ crosslinking to Trm13, non-target tRNAs showed reproducible enrichment that was comparable to canonical targets (Figure S1F and S1G; Tables S6 and S7). Overall, each of the Trm proteins displayed in vivo tRNA binding specificity that was similar but not identical to their known substrate specificities for tRNA methylation. There is precedent for tRNA modifying enzymes to bind with high affinity to tRNAs they do not modify (Keffer-Wilkes et al., 2016; Müller et al., 2013) and our data highlight particular tRNAs for future investigations of the tRNA features that distinguish modification substrates from non-substrates. A few Trm crosslink peaks were observed in nuclear non-coding RNAs (Tables S5-S7). However, it has been noted that certain non-coding RNAs, particularly snoRNAs, appear as frequent hits in eCLIP-Seq experiments even after SMI normalization, and so such peaks should be interpreted with caution (Van Nostrand et al., 2016). Together, these transcriptome-wide crosslinking results show that the known tRNA 2′-O-methyltransferases in yeast interact with diverse RNAs in vivo and have the potential to modify hundreds of additional substrates including mRNAs.

### Transcriptome-wide Mapping of 2′-O-Methyl Ribose with MeTH-seq

To determine whether yeast mRNAs contain 2′-O-methyl ribose (Nm) and define the global landscape of ribose methylation, we developed sequencing of methylation at two prime hydroxyls, MeTH-seq, a high-throughput approach to map Nm sites with single-nucleotide resolution. Nm causes reverse transcriptase (RT) to pause one nucleotide 3′ to the methylated site, a pause that can be selectively enhanced by limiting the availability of deoxyribonucleotides (Maden et al., 1995) or magnesium (Mg^2^+) in the reaction (Figure S2A). We exploited this effect to map the locations of likely Nm sites using next-generation sequencing to identify Mg^2^+-sensitive RT pause sites (Figure 2A; Methods). Different reverse transcriptases and reaction conditions were compared to optimize the sensitivity and specificity of MeTH-seq to identify known 2′-O-methylated sites in rRNA (Figure S2A and S2B). Limiting Mg^2^+ produced slightly better results than limiting dNTPs (Figure S2A). tRNA sites were not considered in this analysis as tRNAs are poorly captured by most RNA-Seq library protocols including ours.

**Figure 2.**
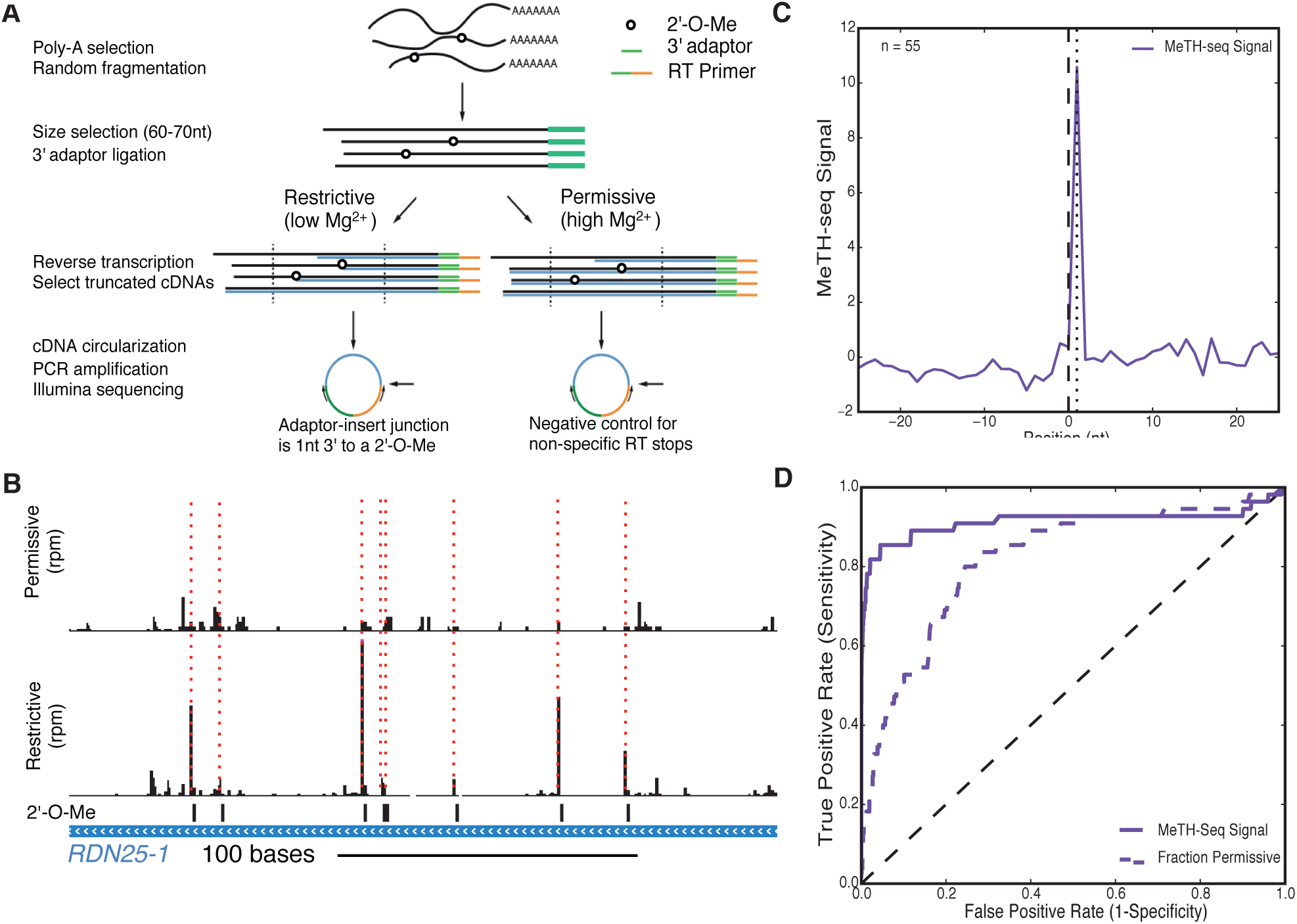
Illumina Sequencing-Based Detection of 2′-O-Methyl Ribose with Single-Nucleotide Resolution. (A)Schematic of MeTH-seq procedure. nt, nucleotides. (B)MeTH-seq reads mapping to a 235 nt region of RDN25-1 (chrXII: 452,800-453,035) containing 8 known Nm sites (black bars). Peaks of Nm-dependent reads are indicated with dashed red lines. (C)Meta plot of MeTH-seq signal for 55 known Nm sites in rRNA. (D)Receiver operating characteristic curve of MeTH-seq signal for all known Nm in rRNA.

MeTH-seq produced clear peaks of reads one nucleotide 3′ of known locations of Nm in rRNA (Figure 2B-2D). Deletion of specific methylation-directing small nucleolar RNAs (snoRNA) eliminated MeTH-seq peaks at their corresponding rRNA target sites (Figure S2C), further confirming the ability of our method to detect Nm sites with single-nucleotide resolution. We used the rRNA data from wild type cells to establish stringent criteria for de novo Nm identification from MeTH-seq data (Methods). 33 out of 55 annotated Nm sites in rRNA were called under these criteria (Table S8; annotations from (Piekna-Przybylska et al., 2007) with the addition of Gm562 in 18S identified by mass spectrometry (Yang et al., 2015). As expected, the MeTH-seq peak caller missed some Nm sites immediately 5′ of other Nm sites, such as Gm805/Am807 in 25S, due to ‘shadowing’ from the downstream site (Table S8). MeTH-seq profiling of wild type yeast ribosomes revealed two likely Nm misannotations: instead of 18S Gm1428 and 25S Am1449, MeTH-seq produced clear signals one nucleotide 5′ to the annotated sites, at Am1427 and Um1448 respectively (Figure S2D). Inspection of previously published primer extension gels supports these changes to the annotations, while a third potential correction (Cm1639 to Gm1638) could not be confirmed or ruled out (gel images available at http://lowelab.ucsc.edu/snoRNAdb/Sc/Sc-snos-bysite.html). After these corrections, Nm sites in rRNA were called with an observed false positive rate of 0.35% (Table S8; Methods). The false positive peaks at Um795 in the 18S and Am816 in the 25S rRNA can be attributed to ‘stuttering’ of RT at Am796 and Am817 (Motorin et al., 2007). A strong Mg^2^+-sensitive RT pause site was observed at 18S U1191, which contains a complex RNA modification, m1acp3Y. A few false positive peaks were found within broader regions of frequent RT pauses that may be due to stable RNA structure. The remaining false positive peaks were qualitatively and quantitatively indistinguishable from peaks at known Nm (Figure S2E and data not shown; Table S8). Importantly, none of the false positive peaks was affected by deletion of snoRNAs (snR72-78). Thus, in subsequent analysis, we required genetic dependence on a 2′-O-methyltransferase to confidently classify a MeTH-seq peak as an Nm site.

Peak heights in rRNA were reproducible between replicates but varied 10-fold between Nm sites whereas the level of methylation likely differs by no more than 2-fold based on mass spectrometry (Figure S2F (Yang et al., 2015). Sequence context is thought to affect the extent of RT pausing at 2′-O-methylated nucleotides (Motorin et al., 2007), and capture biases during RNA-seq library preparation are well known (Raabe et al., 2014). In particular, our approach likely underestimates methylation at NmC and NmG sites (Figure S2G), which may be due to known CircLigase sequence preferences (Lamm et al., 2011). Thus, MeTH-seq peak heights cannot be meaningfully compared between different Nm sites but may be used for relative quantitation of methylation at a given site under different cellular conditions or genetic backgrounds.

### Thousands of Candidate Sites for Regulated 2′-O-Methyl Ribose within mRNAs

Having demonstrated the suitability of MeTH-seq for de novo discovery of Nm sites, we performed MeTH-seq on poly(A)-selected yeast mRNAs during exponential growth in rich medium (A_600 nm_ = 1.0). Given that the stoichiometry of methylation at known rRNA sites is near 100% (Yang et al., 2015), we reasoned that requiring a peak height ≥4 would miss partially methylated sites. The minimum peak height was therefore set as 2.0 for subsequent analyses of potential mRNA 2′-O-methylation. 6,734 sites in 1,947 mRNAs showed a MeTH-seq peak height ≥2.0 in at least 13 out of 16 independent experiments (Figure 3A and 3B; Table S9). However, because we expect more false positives with lower peak heights (Table S10), we subsequently imposed the additional requirement that a peak be genetically dependent on a 2′-O-methyltransferase to be called an Nm site. Relaxing the criteria for Nm identification further, by reducing the requirements for minimum peak height or number of replicates, identified thousands of additional candidate sites (data not shown). Overall, global MeTH-seq profiling suggests Nm may be a prevalent modification in mRNA as well as non-coding RNA, a view supported by genetic evidence as described below.

**Figure 3.**
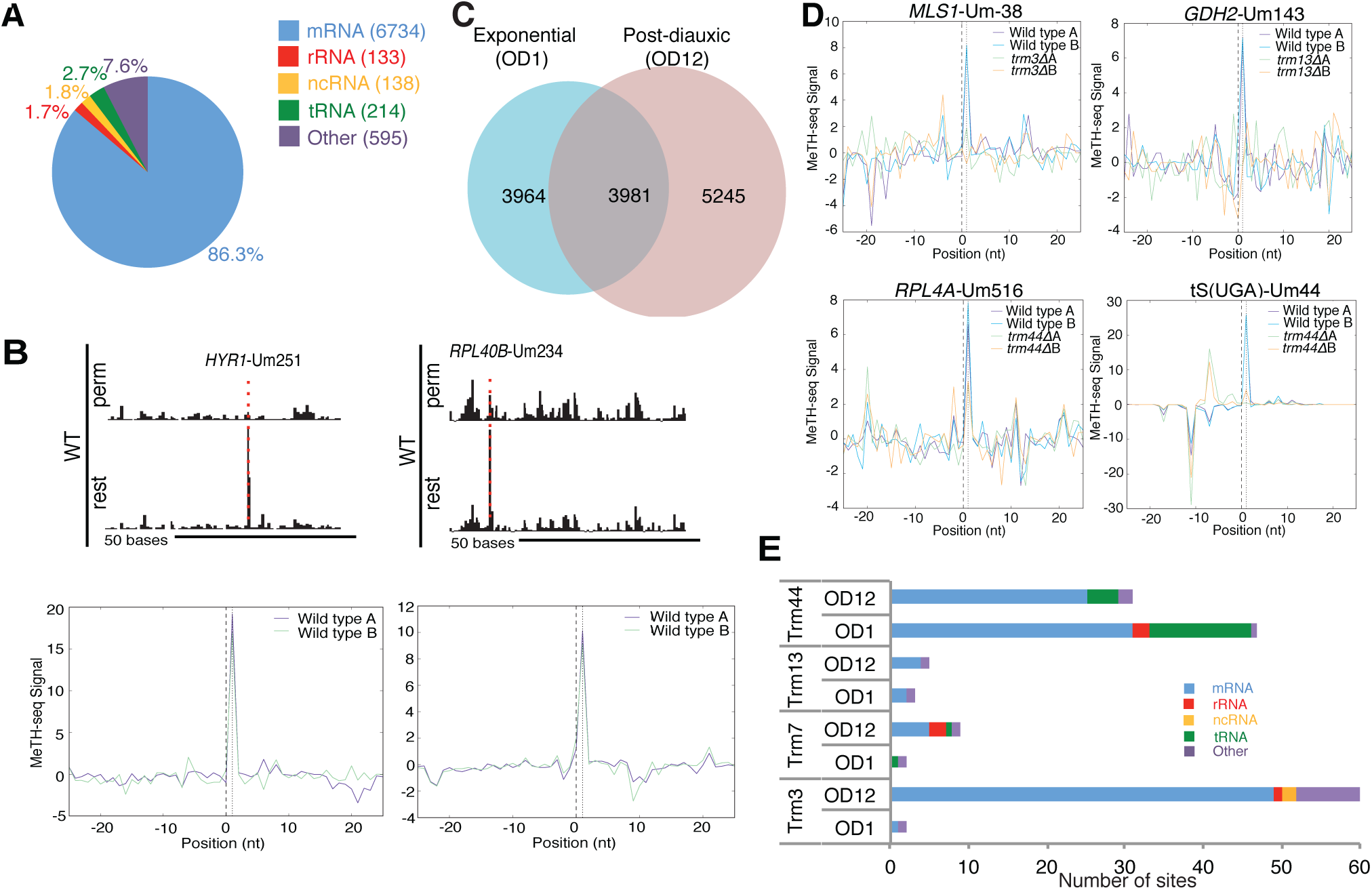
2′-O-Methylation of mRNAs. (A)Distribution of reproducible MeTH-seq peaks between RNA types in exponentially growing yeast. Shown are sites with a peak height ≥2.0 in at least 13 out of 16 independent experiments. (B)Example MeTH-seq peaks in mRNA. Top: genome browser views of reads from restrictive (rest) and permissive (perm) RT conditions. Bottom: MeTH-seq signal plots. (C)Overlap between MeTH-seq peaks identified in exponential (OD1) and post-diauxic (OD12) growth. OD12 sites had peak heights ≥2.0 in at least 16 out of 20 independent experiments. (D)MeTH-seq signal plots of TRM-dependent Nm sites. (E)Summary of Nm sites identified by MeTH-seq as TRM-dependent.

Pseudouridine and m^6^A are dynamically regulated mRNA modifications that respond to nutrient availability and other stresses in yeast (Carlile et al., 2014; Schwartz et al., 2013, 2014). To investigate the possibility of regulated Nm sites, we compared MeTH-seq profiles from exponential and post-diauxic phases of growth, two growth states that differ substantially in gene expression and metabolic activity (Figure S3A and S3B). De novo Nm discovery in post-diauxic cultures (A_600 nm_ = 12) identified 7,900 candidate sites in mRNAs with a MeTH-seq peak ≥2.0 in at least 16 out of 20 independent libraries (Table S11), of which 3,981 were also identified in log phase cultures (Figures 3C).Differences in mRNA abundance do not explain condition-dependent detection of most Nm sites (Figure S3C). Together, these MeTH-seq data suggest extensive 2′-O-methylation of mRNA targets that can be regulated in response to environmental changes.

### Identification of Trm-Dependent 2′-O-Methyl Ribose Sites in mRNAs

The widespread distribution of MeTH-seq peaks together with the evidence that Trm methyltransferases interact with hundreds of yeast mRNAs in vivo suggests that mRNAs could be heavily decorated with 2′-O-methyl ribose. However, the unexplained MeTH-seq peaks (Mg^2^+-sensitive RT pause sites) in rRNA indicates the potential for detection of RNA features or modifications other than Nm in mRNAs. We therefore sought genetic evidence that mRNAs are modified by 2′-O-methyltransferases at the sites identified by MeTH-seq.

We profiled deletion strains lacking each of the four tRNA 2′-O-methyltransferases and identified high-confidence Trm-dependent Nm sites based on reproducible loss of MeTH-seq signal in independent biological replicates of *TRM* deletion mutants (Methods). As expected, deletion of *TRM44* (*trm44Δ*) eliminated signal from Um44 in tRNA^Ser^ (Figure 3D). *TRM13*-dependent methylation of the acceptor stems of tRNA^Gly^, tRNA^His^, and tRNA^Pro^ could not be assessed because the target nucleotides were too close to the 5′ ends of the tRNAs. Likewise, known Trm7 tRNA target sites were not called as MeTH-seq peaks, likely due to the presence of RT-blocking m1A modifications 3′ to Nm32 and Nm34. However, these technical limitations should not affect genetic assignment to a particular 2′-O-methyltransferase of sites already identified by MeTH-seq.

MeTH-seq profiling of *trmΔ* strains identified new RNA methylation targets for each of the Trm proteins in log phase and post-diauxic cells. The largest total number of mRNA candidate Nm sites were assigned to Trm44 (Figures 3D and 3E; Table S12), which is consistent with the presumed greater abundance of Trm44 in poly(A)^+^ RNPs (Beckmann et al., 2015). The relatively low number of *TRM3*-dependent sites, particularly in log phase, was unexpected given the widespread mRNA association observed by eCLIP (Figure 1H). This may reflect technical limitations of the MeTH-seq approach or binding interactions that do not result in modification (Discussion). Overall, our data show that, like the Pus proteins, yeast Trm proteins target specific mRNAs for modification. Thus, ‘moonlighting’ activity towards mRNAs may be a common characteristic among tRNA modifying enzymes.

### Conserved rRNA 2′-O-Methyltransferase Spb1 Targets mRNAs

The majority of candidate Nm peaks were unaffected by deletion of TRMs, suggesting additional 2′-O-methyltransferases may have a role in modifying mRNA. We considered each of the four yeast proteins with validated rRNA 2′-O-methylation target sites: Nop1, Spb1, Mrm1 and Mrm2. Nop1 selects its 18S and 25S rRNA targets through base pairing with a C/D snoRNA guide, which allows computational prediction of potential target sites. All mRNA candidate Nm sites from OD1 and OD12 were examined for possible base pairing with one of 48 snoRNAs followed by filtering for correct positioning of the Nm site at +5 with respect to the snoRNA D box (Methods) (Kiss-László et al., 1996; Nicoloso et al., 1996). Sites positioned at +6 were also included as they are consistent with known snoRNA-directed methylation (e.g. 25S Cm650 targeted by snR18 and Gm1450 targeted by snR24 (Piekna-Przybylska et al., 2007)). Altogether, 10 novel Nm sites were identified as plausible targets of 7 snoRNAs (Figures 4A and 4B; Table S13), suggesting some limited interactions of yeast C/D snoRNAs outside their canonical rRNA targets.

**Figure 4.**
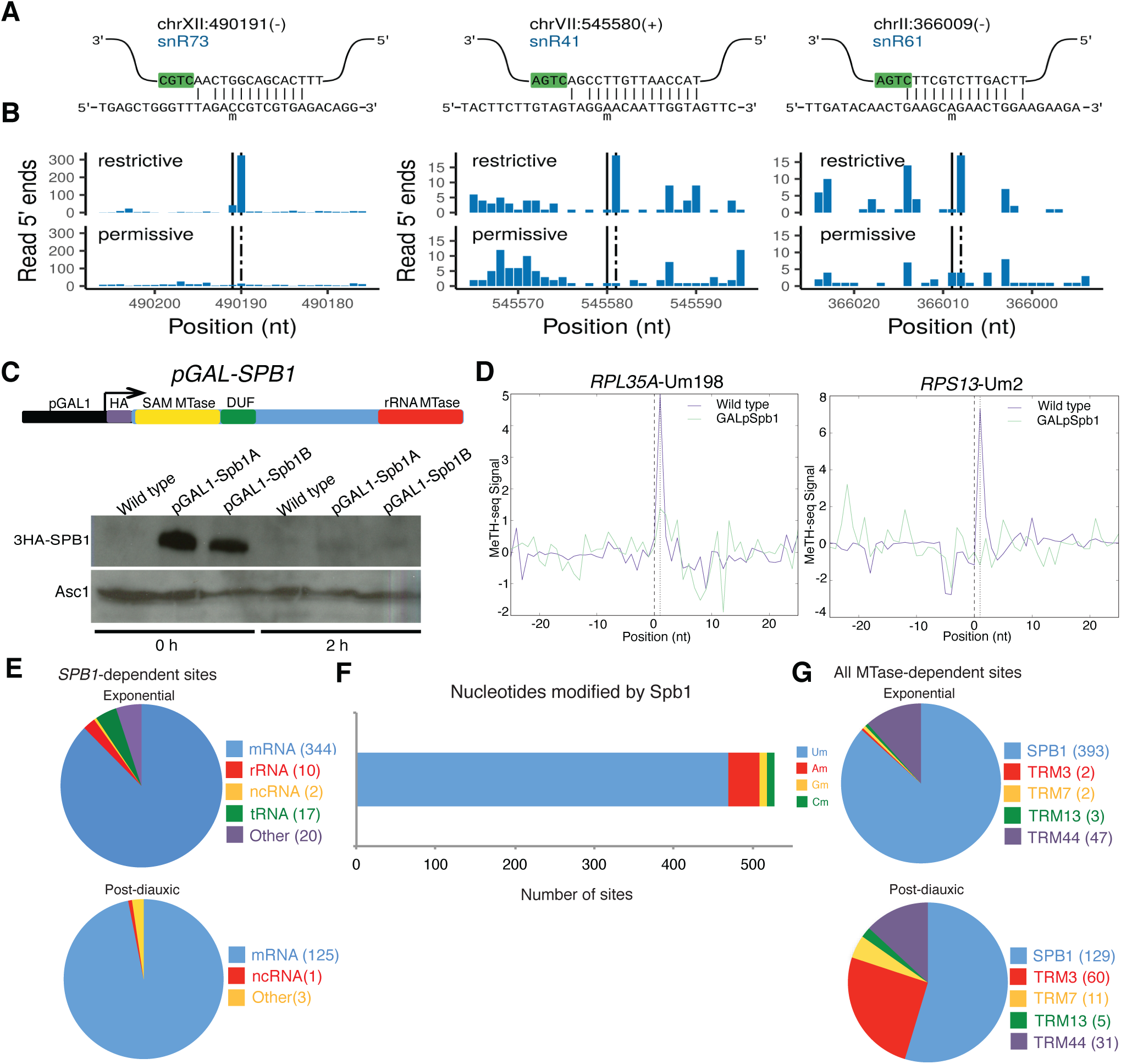
mRNA Methylation by Spb1 Is Widespread. (A)Predicted base pairing between C/D snoRNAs and Nm sites in mRNAs is comparable to canonical snoRNA target sites in rRNA. (B)MeTH-seq reads at putative snoRNA-dependent Nm sites. (C)Depletion of Spb1 by promoter shutoff. Top: The essential *SPB1* gene was placed under control of a repressible GAL1promoter (GALpSPB1) and grown in glucose to inhibit Spb1 expression. Bottom: Tagged Spb1 protein was barely detectable by Western blot after 2 hours. Asc1 loading control was detected as described (Coyle et al., 2009). (D)MeTH-seq signal plots of *SPB1*-dependent Nm sites. (E)Distribution of *SPB1*-dependent Nm sites between RNA types. (F)Distribution of *SPB1*-dependent Nm sites between nucleotides. (G)Methyltransferase dependence of Nm sites in mRNA.

We selected Spb1 for further characterization of the potential for mRNA methylation by rRNA modifying enzymes. Spb1 localizes to the nucleoplasm in addition to the nucleolus and thus has the potential to interact with nuclear mRNA (Kressler et al., 1999). In contrast, Mrm1 and Mrm2 localize to mitochondria where they methylate 21S mitochondrial rRNA (Breker et al., 2013; Pintard et al., 2002b; Sirum-Connolly and Mason, 1993). In addition to its canonical methylation target, 25S rRNA, we found by eCLIP that Spb1 crosslinked to numerous mRNAs (Figure S4A; Table S14). To identify *SPB1*-dependent sites of 2′-O-methylation, the essential *SPB1* gene was placed under control of a repressible *GAL1* promoter (GALpSPB1) and grown in glucose to inhibit *SPB1* expression prior to global MeTH-seq analysis in log phase (OD1) and post-diauxic cells (OD12) (Figure 4C; Methods). As expected, prolonged cell culture in the absence of Spb1 reduced MeTH-seq signal at the known 25S target site, Gm2922, with a peak height of 0.9-1.3 in Spb1-depleted cells compared to 2.6±0.7 across all *SPB1*^+^ libraries (Bonnerot et al., 2003; Lapeyre and Purushothaman, 2004). In addition, signal at 393 (OD1) and 129 (OD12) Nm sites was reproducibly diminished following depletion of Spb1 (Figure 4D; Tables S15 and S16). 87.5% of *SPB1*-dependent Nm sites mapped to mRNAs of which 88.9% were Um (Figures 4E and 4F), which is consistent with the known ability of Spb1 to methylate uridine (Bonnerot et al., 2003; Lapeyre and Purushothaman, 2004). mRNA signal was strongly reduced following Spb1 depletion with 181/393 OD1 and 114/129 OD12 sites decreased ≥4-fold (Tables S15 and S16). In contrast, rRNA sites affected by Spb1 depletion showed modest though reproducible reductions only in log phase, e.g. 25S Um1888 was reduced by ∼20% (Figure S4B-S4D; Table S14). Given that Um1888 and similarly affected rRNA sites are known targets of C/D snoRNAs, these reductions in methylation are likely to be indirect effects of Spb1 depletion although we cannot exclude redundant targeting by Spb1 and snoRNAs (Bonnerot et al., 2003; Lapeyre and Purushothaman, 2004). Of the five tested 2′-O-methyltransferase, our data identify Spb1 as the predominant enzyme acting on mRNAs (Figure 4G).

### Distribution of 2′-O-Methyl Ribose within mRNA Coding Sequences

*SPB1*-dependent MeTH-seq peaks were found in all regions of mRNAs with significant enrichment of sites in CDS (Bonferroni corrected Fisher’s exact test *p* < 8.1e-9) and depletion of sites from 3′ UTRs (*p* < 1.4e-6) (Figures 5A and 5B). Within coding regions Nm was detected in 41 out of 61 sense codons and 1 out of 3 stop codons with enrichment of specific A, I, L, V and M codons (Figure 5C, 5D and S5A; Methods). 18/33 methylated M codons were AUG initiation codons, accounting for the higher density of Nm at the start of CDS regions (Figure 5B). Nm sites were more frequently observed in the 3^rd^ position of sense codons, a distribution of codon positions that differed significantly from a uniform background (Chi Square p<2.2e-16) (Figure 5E). Based on reported position-specific translational effects of Nm on translation in *E. coli* lysates (Hoernes et al., 2015), we analyzed ribosome footprint profiling data from wild type yeast or *dom34Δ* mutants defective in No-Go Decay (Guydosh and Green, 2014) but found no evidence of translation pausing at Nm sites either within ORF bodies or at start codons (Figure S5B-S5D; Methods).

**Figure 5.**
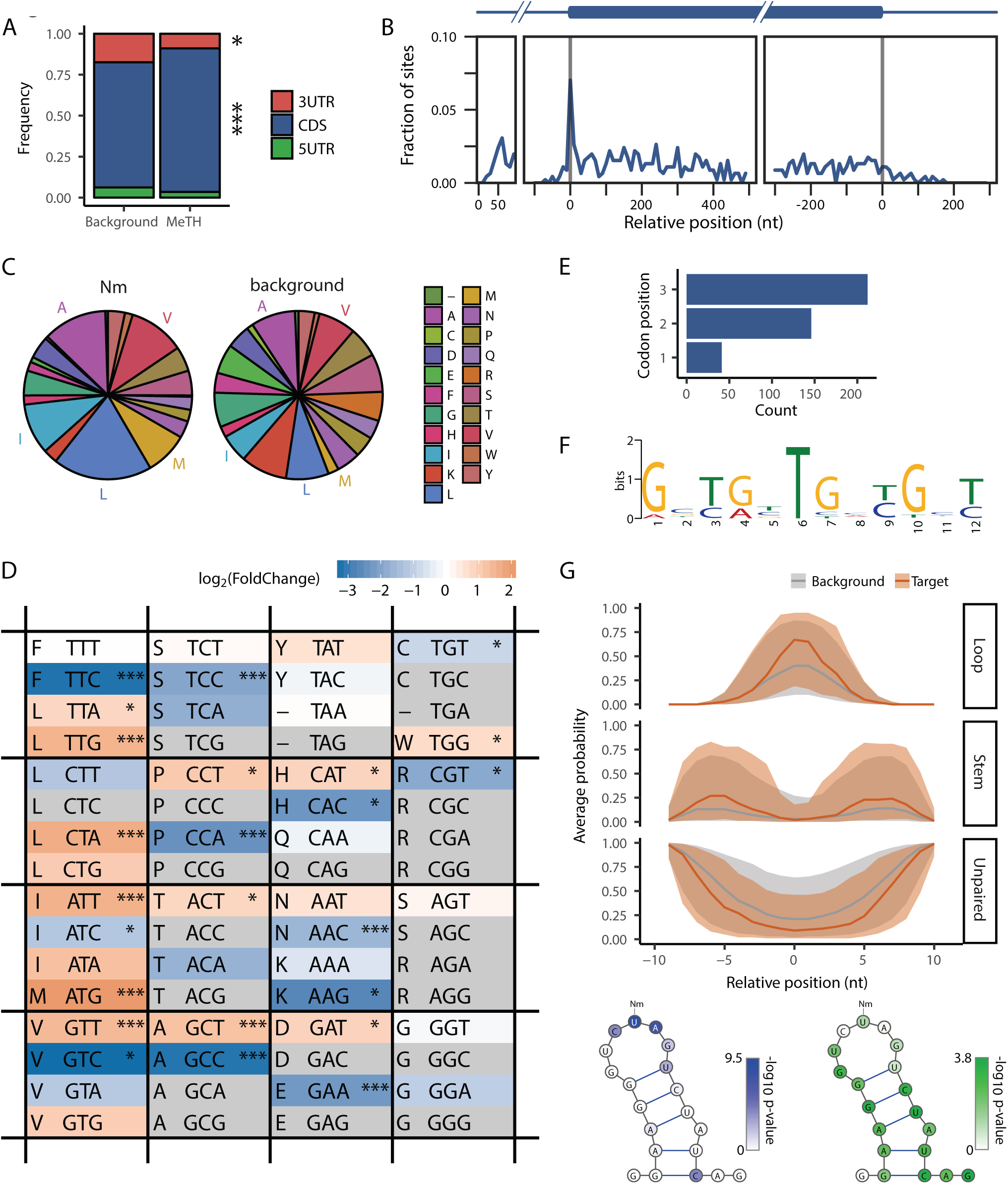
Features of mRNA Target Sites of Spb1. (A)*SPB1*-dependent Nm sites are enriched in CDS (\*\*\**p* < 8.1e-9) and depleted from 3′ UTRs (\**p* < 1.4e-6). Bonferroni corrected *p*-values from Fisher’s exact test. (B)Distribution of *SPB1*-dependent Nm sites with respect to transcription start sites, translation start and stop sites. Numbers indicate distance in nucleotides from landmark. (C-D) Amino acid and codon frequencies among *SPB1*-dependent Nm sites compared to the background of all codons with adequate read coverage for MeTH-seq analysis (Methods). (C) Amino acids A, I, L, M and V are significantly overrepresented. (D) Codon table colors reflect the fold increase or decrease in Nm site frequency compared to background. Codons that did not contain any identified sites are indicated in grey. * *p* < 0.01, *** *p* < 0.0001, Student’s T test. (E)Bias in codon positions methylated by Spb1 **(Chi Square p<2.2e-16)**. (F)Web logo of top motif enriched in Spb1 target sites. E value = 1.7e-40 (Methods). (G)Structural context of Spb1 sites. Each panel plots the probability that a position in Spb1 targets (orange) or the background set (grey) assumes a loop, a stem, or an otherwise unpaired position. Top and bottom edges of the ribbons indicate 75^th^ and 25^th^ percentiles respectively. *p-*values for each position (KS-test) are indicated in the structure diagrams below—left: loops, right: stems.

To explore potential mechanisms underlying mRNA recognition by Spb1, we examined its targets for enriched RNA motifs using MEME (Bailey et al., 2015). A motif rich in UG dinucleotides was notably enriched within a 50-nucleotide window surrounding *SPB1*-dependent Nm sites in mRNA (E value = 1.7e-40; Figure 5F: Methods). This motif was present in 71/409 mRNA target sites while other significantly enriched motifs occurred much less frequently (Figure S5E). The UGNUGN motif was located at variable distances from the modified site (Figure S5F), which was almost invariantly U (Figure 4F). Non-coding RNA targets of FtsJ methyltransferases, including Spb1, are modified on unpaired nucleotides within stem-loop structures (Guy and Phizicky, 2014), but the importance of this structure for Spb1 activity is unknown (Bonnerot et al., 2003; Lapeyre and Purushothaman, 2004). Using RNAsubopt to identify the probable structures of Spb1 targets (Methods), we found that the modified sites were significantly likelier to be in the loop of a hairpin than a background set of sites. Similarly, positions >= 4 nt upstream or downstream of the site were significantly likelier to be involved in base pairing (Figures 5G and S5G). The basis for site-specific methylation by Spb1 remains to be determined and may involve distinct co-factors for different sites as shown for the related methyltransferase Trm7 (Guy and Phizicky, 2014).

### Spb1 Methylates and Maintains Normal Levels of mRNAs Required for Ribosome Assembly

To gain insight into the biological roles of mRNA methylation by Spb1, gene ontology (GO) enrichment analysis was performed. GO terms related to ribosomes, ribosome biogenesis, and translation were almost exclusively over-represented (Figure 6A; Table S17). These related GO term enrichments were driven by the presence of *SPB1*-dependent Nm sites in 97/139 (69%) mRNAs encoding cytoplasmic ribosomal proteins (RP mRNA). “Translational elongation” was also significantly enriched due to methylation of mRNAs encoding elongation factors EF-1 alpha (*TEF1/2*), EF-1B (*TEF4*), EF-2 (*EFT1*), and eIF-5A (*HYP2*) in addition to proteins of the ribosomal stalk (*RPP1A/B* and *RPP2A/B)* that promote recruitment of elongation factors to ribosomes (Gonzalo and Reboud, 2003). Thus, Spb1 targets a coherent regulon of factors required for ribosome assembly and function that includes both its canonical pre-rRNA substrate and numerous RP mRNAs.

**Figure 6.**
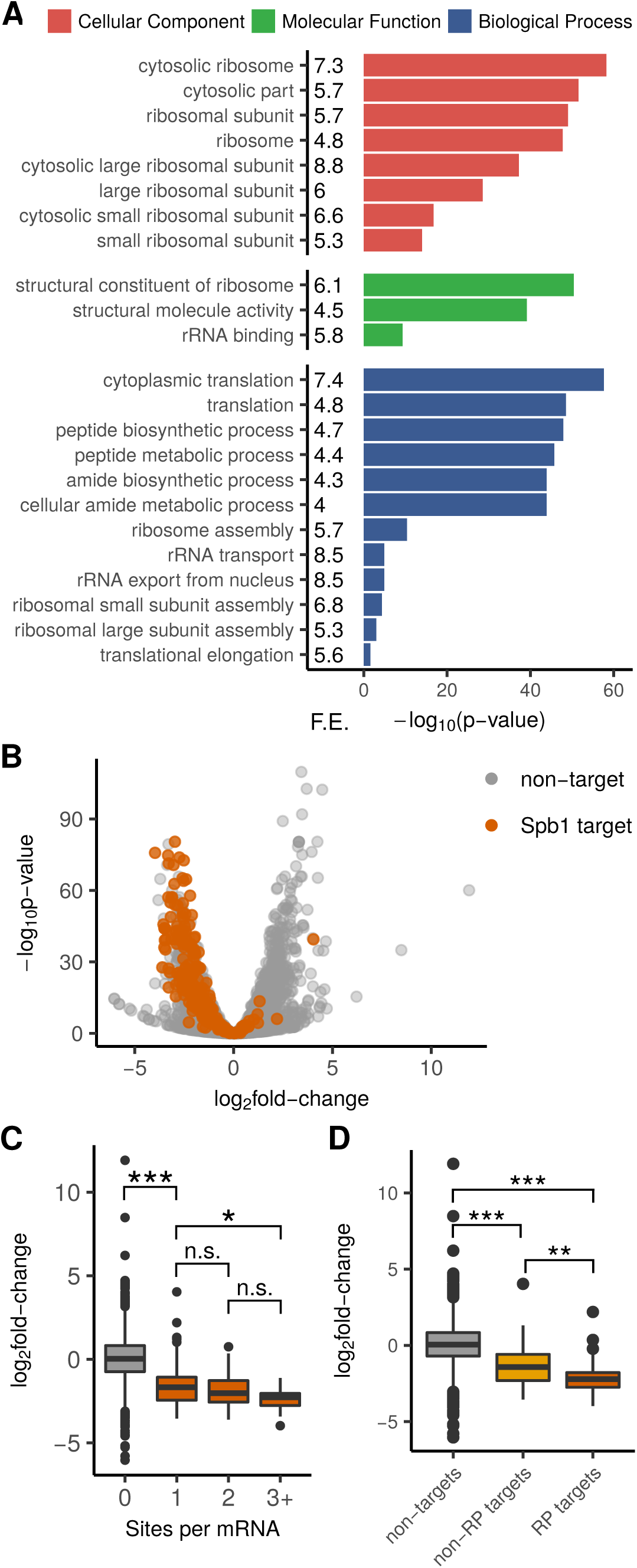
Spb1 Methylates and Maintains Normal Levels of mRNAs Necessary for Ribosome Biogenesis. (A)Log10 *p*-values for GO term enrichments among Spb1 target genes. Displayed categories were enriched ≥4-fold compared to the background of all genes with adequate read coverage for MeTH-seq analysis in Spb1-depleted cells. GO enrichment *p*-values were obtained using PANTHER v13.0. (B)Volcano plot of RNA-seq data shows reduced levels of Spb1 target mRNAs in Spb1-depleted cells. (C)mRNAs with multiple *SPB1*-dependent Nm sites show greater reductions in Spb1-depleted cells. KS test * p<0.0166, *** p<2.2e-16. (D)Spb1 target mRNAs encoding ribosomal proteins showed the largest reductions. KS test *p* < 4.31e-08 For RNA-seq analysis, n=2 for Spb1-depleted cells and n=6 for wild type control cells.

What might be the function of mRNA modification by Spb1? To determine whether loss of Spb1 affects the abundance of its target RNAs we performed RNA-seq on Spb1-depleted and control cells (Methods). In wild type cells, Spb1 target mRNAs were highly expressed with a median TPM (transcripts per million) of 608.5 compared to 15.5 TPM for all genes with adequate read coverage for MeTH-seq analysis (KS test *p* < 2.2e-16; Figure S6A). Depletion of Spb1 lead to significantly decreased abundance of most targets, with somewhat larger changes for mRNAs containing multiple Nm sites (Figure 6B and 6C). RP target mRNAs showed the largest reductions (Figure 6D). Although some of this difference may stem from global effects of Spb1 depletion on RP expression (Figure S6B), target RP mRNAs were significantly reduced compared to non-target RP mRNAs (KS test, *p* < 5.9e-15; Figure S6C). This reduction in mRNA target levels following methyltransferase depletion was specific to Spb1: Trm44 targets increased slightly in *trm44Δ* (KS test, *p* < 5.7e-7; Figure S6D and targets of the other methyltransferases were not significantly affected by the corresponding knockout (data not shown). These data show that *SPB1* is required to maintain the levels of its target mRNAs and suggest Spb1-dependent mRNA methylation may promote mRNA stability. Because a majority of cytoplasmic ribosomal protein mRNAs are targets of *SPB1*-dependent methylation, the role of *SPB1* in ribosome biogenesis is likely to be broader than previously appreciated (Discussion).

## DISCUSSION

Here we identify Spb1 as a conserved methyltransferase that modifies hundreds of yeast mRNAs with 2′-O-methyl ribose (Nm). We further show that *SPB1* expression is required to maintain normal levels of Spb1 target RNAs including most mRNAs encoding ribosomal proteins. Our results expand the known epitranscriptome and establish methods to discover Nm sites transcriptome-wide and illuminate the mechanisms underlying regulation of this novel mRNA modification that is likely to be conserved throughout evolution.

### Detection of 2′-O-methyl Ribose – Methods and Limitations

Three next-generation sequencing methods for Nm profiling have recently been described. Our MeTH-seq library preparation is very similar to the 2OMe-seq method used to identify new Nm sites in mammalian ribosomes (Incarnato et al., 2017), but limits Mg^2^+ rather than dNTPs during reverse transcription, which we found to modestly improve selectivity for Nm. The RiboMethSeq approach exploits the resistance of 2′-O-methylated sites to hydrolysis (Krogh et al., 2016; Marchand et al., 2016). An advantage of this method is the possibility to determine the absolute stoichiometry of modification at specific positions by quantifying the depletion of 3′ and 5′ read ends. However, very high read depth is required to observe depletion of reads at an Nm site making it costly for transcriptome-wide profiling. The Nm-seq method exploits the differential reactivity of Nm to periodate oxidation for selective capture of RNA fragments with Nm 3′ ends (Dai et al., 2017). Pre-enrichment of Nm-containing RNA fragments (e.g. with an antibody) prior to MeTH-seq, 2OMe-seq or RiboMethSeq would allow comprehensive Nm discovery, with single-nucleotide resolution, at much lower sequencing depth. However, enrichment of modified RNA fragments prior to sequencing, which is standard in m6A and m1A profiling studies, may result in capture of RNAs that are modified at very low stoichiometry and have uncertain physiological relevance. In contrast, as implemented here, our approach requires RT to pause on a substantial fraction of total RNA molecules to detect a peak for a given Nm site.

MeTH-seq may not capture all 2′-O-methylated sites. It is notable that MeTH-seq identified relatively few *TRM3*-dependent Nm sites despite the widespread mRNA association of Trm3 observed by eCLIP. Remarkably, >96% of high-confidence binding sites identified in the Trm3 eCLIP libraries mapped to mRNAs. Altogether, we identified 1611 stringent peaks in 975 mRNAs in addition to 29 stringent peaks that mapped to 19 tRNAs. Such pervasive mRNA binding by Trm3 was unexpected, and the basis for it is not immediately apparent. Trm3 does not have known RNA-binding domains outside the conserved methyltransferase catalytic domain. Furthermore, Trm3 is not more highly expressed than the other tRNA 2′-O-methyltransferases (Ghaemmaghami et al., 2003; Kulak et al., 2014), and all four HPM-tagged Trm proteins were immunoprecipitated with similar efficiency. Given that Trm3 methylates its canonical tRNA target sites in the context GmG, this discrepancy between detection of binding and modification may reflect false negatives due to sequence capture bias in the MeTH-seq protocol.Specifically, truncated cDNAs ending in C are inefficiently circularized by CircLigase (Lamm et al., 2011). In addition, the true methylation targets may interact with Trm3 very transiently and therefore be poorly captured by CLIP. Alternatively, Trm3 crosslinking may enrich for deadenylated mRNAs that were not examined by our profiling of Nm sites in poly(A)^+^ RNA. There is also precedent for tRNA modifying enzymes to bind with high affinity to non-substrate RNAs, perhaps to affect RNA folding (Keffer-Wilkes et al., 2016; Müller et al., 2013).

The majority of highly reproducible MeTH-seq peaks (those present in at least 13 independent samples) identified here were not affected by depletion of known 2′-O-methyltransferases. As the TRM methyltransferases and Spb1 do not modify one another’s non-coding RNA targets in vivo, we consider widespread redundant targeting of mRNA sites by these enzymes to be unlikely. The mitochondrial enzymes Mrm1 and Mrm2, whose modification targets were not determined in our study, conceivably methylate cytoplasmic mRNAs either en route to the mitochondria or from a cytoplasmic pool of protein (Breker et al., 2013). Consistent with this possibility, it is interesting that the ortholog of Mrm1 crosslinks to poly(A)^+^ RNA in human cells (Baltz et al., 2012; Castello et al., 2012). In addition, the *S. cerevisiae* genome encodes multiple predicted RNA methyltransferases of unknown activities (Wlodarski et al., 2011), which potentially include additional mRNA 2′-O-methyltransferases.

However, although other known or putative RNA methyltransferases may install Nm in mRNA, it is likely that other RNA features, such as structure, can give rise to reproducible Mg^2^+-sensitive RT pause sites in the absence of 2′-O-methylation. In any sequencing library preparation there is the potential for robust artefacts, such as mispriming during reverse transcription (Gillen et al., 2016), which may explain the enrichment of a primer sequence motif among candidate Nm sites proposed from Nmseq data (Dai et al., 2017). Therefore, we do not interpret MeTH-seq peaks as Nm sites in the absence of genetic validation. The remaining potential mRNA 2′-O-methyltransferases need to be tested before the complete Nm landscape can be known in yeast.

### Conservation of mRNA 2′-O-methylation

While our study was being finalized, He and colleagues reported the presence of Nm in mRNA from human cells based on mass spectrometry and sequencing-based approaches, although they did not identify the methyltransferases responsible (Dai et al., 2017). Each of the mRNA modifying yeast methyltransferases characterized here is conserved to man. Notably, FTSJ3, which is the human ortholog of Spb1, here identified as a predominant mRNA methyltransferase in yeast, has been found in the poly(A)^+^ mRNP interactomes of multiple human cell lines (Baltz et al., 2012; Beckmann et al., 2015; Castello et al., 2012). Proteomic characterization of FTSJ3 co-immunoprecipitates identified 10 hnRNP proteins providing further evidence that FTSJ3, like Spb1, may interact with nuclear mRNA (Simabuco et al., 2012). Our analysis also suggests some mRNAs are targeted by the snoRNA-directed methyltransferase Nop1, whose human counterpart fibrillarin was likewise found to crosslink to poly(A)^+^ human RNA. Although the reported locations of Nm in human mRNA have been called into question based on the apparent mispriming artefact described above, mass spec analysis establishes 2′-O-methylated nucleotides as abundant in purified bulk human mRNA (Dai et al., 2017).Furthermore, Um was by far the most abundant with a molar ratio of 0.15% for Um/U, which is consistent with the predominance of Um among genetically validated Spb1-dependent Nm sites. Thus, we conclude that 2′-O-methyl ribose is conserved as an mRNA modification in diverse eukaryotes and suggest that the mRNA methylation activity of Spb1/FTSJ3 in particular is likely to be conserved.

### Functional consequences of Nm in mRNA

We found that depletion of Spb1 led to substantial reductions in the levels of its mRNA targets, which suggests endogenous Nm could promote mRNA stability. 2′-O-methylated nucleotides are widely used in synthetic small interfering RNA (siRNA) due to their resistance to degradation by various nucleases (Czauderna et al., 2003; Sproat et al., 1989). However, it is unclear how the presence of sparse Nm residues within coding sequences would stabilize an mRNA against exonucleolytic decay from the 5′ or 3′ end of the transcript. If Nm increases steady-state mRNA levels by conferring resistance to endonucleolytic cleavage at the modified site, this uncharacterized decay pathway must play a sizable role in yeast mRNA homeostasis. Alternatively, Nm may recruit a protein factor that promotes mRNA stabilization. Such a mechanism would be opposite to mRNA destabilization by m6A, which occurs through direct binding of YTHD family proteins to modified mRNA and subsequent recruitment of decay factors (Ke et al., 2017; Wang et al., 2014). Any Nm ‘reader’ likely recognizes the modified nucleotide in a broader sequence or structural context; recognition of additional RNA features could explain why loss of Nm affects the abundance of Spb1 targets specifically whereas loss of Nm did not decrease the levels of mRNAs targeted by other methyltransferases.

Consistent with this possibility, the presence of Nm has been found to affect interactions with several RNA binding proteins in both natural and artificial contexts (Devarkar et al., 2016; Lacoux et al., 2012; Lavoie and Abou Elela, 2008; Simon et al., 2011; Tian et al., 2011). With the exception of PAZ domains binding the 2′-O-methylated 3′ ends of small RNAs, most characterized RNA binding protein interactions with 2′-O-methylated RNAs show reduced protein binding to modified RNA. In contrast, m^6^A is known to promote binding to multiple ‘reader’ proteins to mediate a variety of downstream effects on mRNA metabolism (Roignant and Soller, 2017; Yue et al., 2015). Our findings motivate the search for Nm readers and their functional interactions with mRNA decay pathways.

We anticipated that mRNA 2′-O-methylation might impede translation elongation based on results with synthetic Nm containing mRNAs (Dunlap et al., 1971; Hoernes et al., 2015). We therefore analyzed ribosome footprint profiling data to determine ribosomal A site occupancy at modified codons. No pausing was observed either in wild type cells or in *dom34Δ* mutants that are defective in ribosome stalling-dependent No-Go Decay (Guydosh and Green, 2014) nor did deletion of *DOM34* increase steady-state levels of translating Nm-containing mRNAs. Given that Spb1 methylation targets are among the most highly expressed and efficiently translated mRNAs in growing yeast, it would be surprising if their expression were inhibited at the level of translation elongation.Furthermore, depletion of Spb1 decreased target mRNA levels, which is the opposite of what would be expected for a modification that slowed elongation (Hanson and Coller, 2017). The lack of detectable ribosome pausing at Nm codons in vivo may reflect more robust translation elongation in cells compared to lysates. However, CAmA, which caused a 10-fold reduction in synthesis of full-length protein in *E. coli* lysates (Hoernes et al., 2015), was found infrequently among Nm-containing codons in yeast and the ribosome profiling sequencing depth was not sufficient to assess pausing at this small number of codons in isolation. Thus, we cannot exclude the possibility that specific Nm modified codons could inhibit elongation in cells.

### A Broader Role for Spb1 in Ribosome Biogenesis?

A wealth of genetic and biochemical evidence implicates *SPB1* in ribosome biogenesis, but the essential function of the Spb1 protein in this process remained unclear. *SPB1* was originally characterized as one of seven Suppressors of Poly(A) Binding Protein that affect the levels of 60S ribosomal subunits in yeast (Sachs et al., 1987). Depletion of Spb1, which co-purifies with 66S pre-ribosomal particles, leads to accumulation of rRNA processing intermediates and depletion of mature 25S rRNA (Harnpicharnchai et al., 2001; Kressler et al., 1999). The subsequent discovery that Spb1 methylates 25S rRNA revealed a molecular function for its conserved methyltransferase domain (Bonnerot et al., 2003; Lapeyre and Purushothaman, 2004). Mutating the predicted catalytic aspartate (*spb1-D52A*) abolished the 25S Gm2922 modification and produced a viable but severely growth impaired strain with greatly reduced 60S levels (Lapeyre and Purushothaman, 2004). However, it is remarkable for a single Nm site to be critical for ribosome biogenesis. Indeed, most yeast C/D snoRNAs can be deleted with little effect on growth. Thus, the apparent importance of Spb1-dependent RNA methylation for ribosome production and cell growth was puzzling.

Our work identified hundreds of new RNA targets for 2′-O-methylation by Spb1 that include the majority of mRNAs encoding cytoplasmic ribosomal proteins. Depletion of Spb1 led to significantly reduced levels of its mRNA targets consistent with a positive role for *SPB1* – and presumptively RP mRNA 2′-O-methylation – in ribosomal protein production. Spb1 modification targets included RP mRNAs encoding proteins of both ribosomal subunits, which could contribute to the reduction in 40S observed following longer depletion of Spb1 (Kressler et al., 1999). Of note, although the human ortholog FTSJ3 is also required for ribosome biogenesis, it contributes primarily to 40S production (Morello et al., 2011). It will be important to learn whether FTSJ3 similarly modifies RP mRNAs and determine how mRNA 2′-O-methylation affects RNA levels in human cells.

## ACKNOWLEDGEMENTS

We thank E. Van Nostrand and G. Yeo for eCLIP advice; B. Waldman for help with MeTH-seq feasibility studies; C. Burge, E. Phizicky and members of the Gilbert Lab for discussion. The sequencing was performed at the MIT BioMicro Center under the direction of S. Levine. This work was supported by grants from The American Cancer Society – Robbie Sue Mudd Kidney Cancer Research Scholar Grant (RSG-13-396-01-RMC) and the National Institutes of Health (GM094303, GM081399) to W.V.G. T.M.C. and K.M.B. were supported by fellowships from The American Cancer Society. C.S. was supported by an NSF GRF.

## AUTHOR CONTRIBUTIONS

K.M.B. designed research and performed and analyzed MeTH-seq experiments. T.M.C. developed the MeTH-seq data analysis pipeline. C.S. designed and performed all other computational analyses. W.V.G. conceived the project, supervised research, designed research, and performed eCLIP. K.M.B., C.S. and W.V.G. interpreted the results. W.V.G. wrote the paper with input from all authors.

## TABLES

Table S1: Yeast strains used in this study.

Table S2: Sequencing read summary for eCLIP libraries.

Table S3: Trm44 eCLIP clusters: positions, enrichments and *p*-values.

Table S4: Pus1 eCLIP clusters: positions, enrichments and *p*-values.

Table S5: Trm3 eCLIP clusters: positions, enrichments and *p*-values.

Table S6: Trm7 eCLIP clusters: positions, enrichments and *p*-values.

Table S7: Trm13 eCLIP clusters: positions, enrichments and *p*-values.

Table S8: MeTH-seq peaks in rRNA, peak height ≥ 4.0.

Table S9: All MeTH-seq peaks ≥ 2.0 in log phase (OD1).

Table S10: MeTH-seq peaks in rRNA, 4.0 ≥ peak height ≥ 2.0.

Table S11: All MeTH-seq peaks ≥ 2.0 in post-diauxic phase (OD12).

Table S12: TRM-dependent Nm sites (OD1 and OD12).

Table S13: Summary of predicted snoRNA-dependent Nm sites in mRNA.

Table S14: Spb1 eCLIP clusters: positions, enrichments and *p*-values.

Table S15: *SPB1*-dependent Nm sites in log phase (OD1).

Table S16: *SPB1*-dependent Nm sites in post-diauxic phase (OD12).

Table S17: GO term summary: Spb1 target mRNAs.

## Materials and Methods

### CONTACT FOR REAGENT AND RESOURCE SHARING

Further information and requests may be directed to and will be fulfilled by the Lead Contact, Wendy Gilbert.

### EXPERIMENTAL MODEL AND SUBJECT DETAILS

#### Yeast strains and growth

All yeast strains are *Saccharomyces cerevisiae* BY4741 or BY4742 derivatives (BY4742:wild type (YWG11), BY4741:wild type (YWG506), *TRM44-HPM* (YWG1354), *PUS1-HPM* (YWG1357, YWG1358), *TRM3-HPM* (YWG1348, YWG1349), *TRM7-HPM* (YWG1350, YWG1351), *TRM13-HPM* (YWG1352, YWG1353), *SPB1-FLAG* (YWG1359, YWG1523), *snr72-78Δ* (YWG318, YWG372), *trm3Δ* (YWG1320, YWG1321), *trm7Δ* (YWG1322, YWG1323), *trm13Δ* (YWG1326, YWG1327), *trm44Δ* (YWG1324, YWG1325), *trm732Δ* (YWG1332, YWG1333), *rtt10Δ* (YWG1330, YWG1331), *pGAL1-HA-SPB1* (YWG1522, YWG1523). See Table S1 for complete genotypes. snoRNA deletions, *trm13Δ* and *pGAL-HA-SPB1* strains were made using PCR-based deletion cassettes (Longtine et al., 1998). Other deletion strains were obtained from the Yeast Deletion Collection (Winzeler et al., 1999). His-PrecissionProtease-MYC (HPM) tagged strains were made using PCR-based cassettes (Graumann et al., 2004). All strains, with the exception of *pGAL-HA-SPB1*, were grown at 30°C in YPAD (1% yeast extract, 2% peptone, 0.01% adenine hemisulphate, 2% glucose). Cultures for MeTH-seq were harvested by centrifugation in log phase (A_600nm_≈1.0) or at high density (A_600nm_≈12.0). High density *trm7Δ* cells were grown to A_600nm_≈8.0.

The *pGAL-HA-SPB1* depletion strain was maintained at 30°C in YPARG (YPA with 20% raffinose, 30% galactose). Cells were transferred to YPAD to inhibit expression of *SPB1*. Log phase cells: Cells were grown to an A_600nm_≈0.3, centrifuged and resuspended in YPAD media for an additional 4-6 hours until A_600nm_≈1.0. High density cells: Cells were grown to an A_600nm_≈3.0, centrifuged and resuspended in YPAD media for an additional 8-12 hours until A_600nm_≈12.0.

For ePAR-CLIP assays yeast were grown at 30°C in minimal media with reduced uracil (SC_Ura120_μM) to an A_600nm_=0.4-0.5, supplemented with 4-thiouracil to a final concentration of 500μM and cultured for 3 more hours before harvest as described (Beckmann, 2017).

### METHOD DETAILS

#### Yeast PAR-CLIP

Yeast strains expressing tagged methyltransferases from their endogenous loci were cultured in the presence of 500μM 4-thiouracil. Cells were harvested by centrifugation, resuspended in ice cold water, and UV irradiated on ice in a Stratalinker (365 nm, energy = 7.2J/cm^2^). Crosslinked cells were washed in ice water, flash frozen in liquid N_2_, and stored at −80°C until lysis. Cells were resuspened in 1.5 mL iCLIP lysis buffer (50 mM Tris-HCl pH7.4, 100 mM NaCl, 1% NP-40, 0.1% SDS, 0.5% sodium deoxycholate, 1X protease inhibitor cocktail (Millipore), 400U/mL Murine RNase inhibitor (NEB)) per g of cell pellet and lysed by vortexing with glass beads (Manufacturer). Lysates were clarified by centrifugation, flash frozen in liquid N_2_ and stored at −80°C.

#### eCLIP library preparation

eCLIP libraries were prepared as described (Van Nostrand et al., 2016). Briefly, lysates were diluted in iCLIP lysis buffer to 26 ODU/2 mL and treated with Turbo DNase (Lifetech) and RNase I (Lifetech) for 15min at 22°C before placing on ice. Treated lysates were centrifuged (15,000 × *g* 15min) and added to Protein G magnetic beads (Dynabeads) pre-bound with 10 μg of antibody (anti-Myc or anti-FLAG, Sigma) and rotated for 2.5hr at 4°C. 4% of the binding reaction was saved as input before extensive washing of bead-bound protein-RNA complexes. Bound RNA 3′ ends were dephosphorylated with FastAP (Lifetech) and PNK (NEB) before on-bead ligation of a 3′ RNA linker with T4 Rnl (NEB) at 16°C overnight. After washing, 10% of bound RNA was labeled by PNK with ^32^P ATP for diagnostic gels. The remainder was eluted from the beads by incubation in 1X NuPAGE loading buffer with 0.1M DTT at 70°C for 10min.CLIP eluates and paired inputs were electrophoresed on 4-12% Bis-Tris NuPAGE and transferred to nitrocellulose membranes. Tagged protein MW was determined by western blot of 2% input before CLIP and size-matched input (SMI) membranes were cut, taking a 75kDa region just above the target protein size. RNA was released with proteinase K, extracted with phenol:chloroform:isoamy alcohol, and purified on RNA Clean & Concentrator-5 columns (Zymo). SMI RNA was dephosphorylated, a 3′ linker ligated, and purified on silane beads (Dynabeads). CLIP and SMI RNA was reverse transcribed with SuperScript IV, hydrolyzed with NaOH, neutralized with HCl, and cDNA was purified on silane beads. A 5′ linker was ligated with T4 Rnl (NEB) overnight at 16°C. Linker-ligated cDNA was purified on silane beads before diagnostic PCRs to determine the minimum number of cycles. Final PCR products were gel-purified, precipitated and sequenced on an Illumina HiSeq 2000.

#### RNA isolation

Total yeast RNA was isolated by hot acid phenol extraction followed by isopropanol precipitation as described (Carlile et al., 2015). PolyA+ mRNA was isolated from 10-15mg of yeast total RNA by two rounds of selection on oligo-dT cellulose beads according to the manufacturer’s instructions (NEB).

#### MeTH-seq library preparation

RNA fragmentation was performed in 10mM ZnCl_2_ at 95°C for 55s. RNA fragments were precipitated, dephosphorylated with CIP and PNK, and separated by denaturing (8% urea-TBE) polyacrylamide gel electrophoresis (PAGE). Gel-purified RNA fragments (60-70nt, 70-80nt, 80-90nt) were eluted overnight with rocking in RNA elution buffer (300mM NAOAc pH 5.5, 1mM EDTA, 100 U/ml RNasin (Promega) and ligated to a pre-adenylated adaptor (/5Phos/TGGAATTCTCGGGTGCCAAGG/3ddC/) (IDT) using T4 RNA ligase (NEB) at 22°C for 2.5h, followed by precipitation. Reverse transcription was completed using AMV-RT (Promega) and the reverse transcription primer (/5Phos/GATCGTCGGACTGTAGAACTCTGAACCTGTCGGTGGTCGCCGTATCATT/iS p18/CACTCA/iSp18/GCCTTGGCACCCGAGAATTCCA) (IDT) using the following conditions. RNA and RT primer were denatured and annealed in reverse transcription buffer (50 mM Tris-Cl pH 8.6, 60mM NaCl, 10mM DTT). After annealing, the reverse transcription reaction was split in equal halves for the restrictive and permissive conditions. The restrictive reaction received a final concentration of 2.5mM dNTPs, 6mM MgCl_2_ and 7.1% DMSO, whereas the permissive reaction received a final concentration of 4mM dNTPs, 20mM MgCl_2_ and 7.1% DMSO. Reactions were carried out at 42°C for 1h, prior to degradation of RNA with the addition of NaOH at 85°C. Truncated cDNAs were size-selected and purified on an 8% urea-TBE PAGE gel, followed by elution from gel slices in DNA elution buffer (300mM NaCl, 10mM Tris-HCl, pH 8.0) overnight at room temperature. cDNAs were circularized with circLigase (Epicentre) and amplified by PCR (Phusion; NEB) with the forward primer (AATGATAC GGCGACCACCGA) and a barcoded reverse primer (CAAGCAGAAGAC GGCATACGAGATXXXXXXGTGACTGGAGTTCCTTGGCACCCGAGAATT CCA) (IDT). PCR products were gel-purified, precipitated and sequenced on an Illumina HiSeq 2000.

#### Ribo-seq

Ribosome footprint profiling was performed essentially as described in Thompson et al. (2016), with the following minor modifications. Yeast cells were grown to an A_600nm_≈1.0 or A_600nm_≈12.0, harvested by vacuum filtration (http://bartellab.wi.mit.edu/protocols.html.), lysed using a Cryomill and cycloheximide was added to the lysis buffer (0.1 mg/mL) prior to centrifugation. rRNA was subtracted from mRNA libraries as described (Brar et al., 2012).

### QUANTIFICATION AND STATISTICAL ANALYSES

Most statistical tests were carried out in R, except for the codon enrichment analysis, where the tests were carried out using the scipy package in python. Plots were generated using matplotlib in python, or the ggplot2 package from the tidyverse in R.

#### eCLIP-seq data analysis

Our analysis pipeline is based on the one described by the Yeo lab (Van Nostrand et al, 2016). For full code, see <GitHub repo>. Reads were first demultiplexed by removing the first 10-nt of the forward read and adding them to the sequence name for later use; these 10-nt correspond to the random barcode added as part of the 5′ linker. The adapter sequence was trimmed from raw reads using cutadapt (version 1.7.1), requiring a minimum trimmed read length of 18-nt. Reads were mapped using STAR (version 2.5.1b) to a repeat-masked yeast genome. This genome was generated by masking repetitive regions and non-coding RNA loci (particularly tRNA genes and rRNA repeats) that had at least one identical sequence elsewhere in the genome. To ensure unique copies of these genes, we included a pseudo-chromosome that concatenated single copies of the masked non-coding RNA genes. We used the 10-nt barcodes to collapse PCR duplicates, merging any reads that mapped to the same position and shared the same barcode. Peaks were then called using the CLIPper algorithm. The pulldown and size-matched input libraries were processed separately in this way, and finally peaks from pulldown libraries were normalized to peaks in the input library using a normalization pipeline (Van Nostrand et al 2016).

#### RNA-seq data analysis

RNA-seq data was analyzed with in-house Bash and Python scripts. For a given MeTH-Seq experiment, we used the reads from the permissive library to quantify gene expression. Reads were trimmed using cutadapt, requiring at least 5 nucleotides of overlap between the read and the adapter, and requiring trimmed reads to be at least 18-nt long. Trimmed reads were mapped to yeast mature mRNA sequences (obtained from SGD) and quantified using kallisto (version 0.43.1), using options --single -l 40 –s Differential expression was calculated using DESeq2 (version 1.14.1).

#### Identification of Nm sites

Sites of 2’-O-methylation were identified as previously described for pseudouridine (Stanley et al., 2016) with modifications. MeTH-seq signal was calculated as follows. For each position *i* in a 51 nt window centered at a given genomic position, the fraction of reads in the window whose 5′ ends map to *i* was calculated. MeTH-seq signal is the difference in fractional reads between the restrictive and permissive libraries, multiplied by the window size. For an identified Nm, the reported MeTH-seq signal corresponds to the RT stop position 1 nt 3′ of the site. All genomic positions with coverage of ≥ 0.25 reads per nt in the above described window within annotated features (downloaded from SGD on 9/2/2011) were considered, including 5’ and 3’UTRs identified in (Pelechano et al., 2014). For Nm calling we required a reproducible peak height ≥ 2.0 with reproducibility in at least 13 of 16 log phase libraries and at least 16 of 20 high density libraries. A window size of 51 nt was examined in all libraries and only windows that surpassed the read cutoff of 0.25 reads per nt were considered.

#### Methyltransferase assignment

MeTH-seq peak heights were compared between methyltransferase deletion strains and all libraries from a given growth condition from cells wild type for a given methyltransferase. An Nm site was called genetically dependent on a specific Trm enzyme if the peak heights in both *trmΔ* replicates were 1.5 standard deviations below the mean across all libraries. This threshold was set based on the observed reductions in peak heights at positive control tRNA^Ser^ Um44 sites in *trm44Δ* libraries. Spb1 target sites were called at a more stringent threshold requiring peak heights at least 2.0 standard deviations below the mean in both Spb1-depleted libraries and coverage of ≥0.25 reads per nt in the 51nt window surrounding the Nm site in both Spb1-depleted replicates.

#### MeTH-seq signal plots and ROC curves

MeTH-seq signal plots show the average of the MeTH-seq signal over the 51 nt window (described above) for all Nm included in the analysis. To generate receiver operating characteristic (ROC) curves for a given permissive/restrictive library pair, MeTH-seq signal was calculated for each position within the rRNA. A range of 5000 equally spaced cutoff scores were chosen spanning the range of observed MeTH-seq signal values. At each cutoff score, the true positive and false positive rates were calculated, and plotted.

#### SnoRNA target analysis

In this analysis, we did not attempt to predict potential target sites across the transcriptome; instead, we aimed to identify which of the putative sites Nm sites were consistent with modification by snoRNAs. To do this, we generated a fasta file of the 30-nt sequences surrounding each site, which we then used to generate a BLAST database. We identified short complementary regions between the snoRNAs and the target sequences by running a BLAST search, using all yeast C/D box snoRNAs as queries and the options-task blastn-short-strand minus. We filtered the list of hits based on the mechanism of C/D box targeting: we required that the aligment include the putative Nm position, and that this position were 5 or 6 nt downstream of the position opposite a D-box.

#### Codon enrichment analysis

We calculate background frequencies for each codon in each of the N wild-type libraries used for peak calling at OD1. To calculate the frequency of a codon *i* in a given library, we weigh the expression (RPKM) of each gene by the number of times it contains codon *i*, then sum across these and normalize by the total.

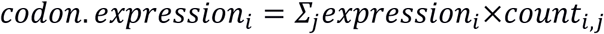

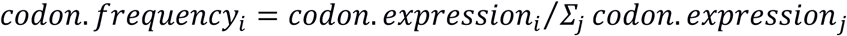

We compare the frequency of each codon among Spb1 target sites to its background distribution using the SciPy implementation of the Student’s T test.

We calculate the reported fold-change as the ratio of the codon frequency in Spb1 sites and the average frequency in the background distribution.

#### Motif detection

MEME (version 4.12.0) was used to find enriched motifs among Spb1-dependent mRNA sites. Detection was carried out on the 50-nt sequences surrounding each of the Spb1-dependent sites, calculating motif enrichment relative to a 2-order background model that summarized trinucleotide frequencies in the set of target sequences. The motif search required that any identified motifs be at least 3-nt and at most 15-nt long. The background model was generated using the fasta-get-markov script that is included with the MEME package, using the following options: -m 2 -rna -norc.

#### Pairing probability distribution

Pairing probabilities of 20-nt regions surrounding Spb1-dependent 2’OMe sites were compared to a background set consisting of randomly selected sites from genes that contained at least one Spb1-dependent site. For each sequence, RNAfold (ViennaRNA package version 2.3.3) was used to obtain a pairing probability matrix. We then summed across the columns of the matrix to calculate the probability that each position is involved in base-pairing. To obtain a background distribution of average pairing probabilities, the following was done for each of 100 iterations: randomly chose 391 sites from the background sets (such that each background set is as large as the target set), then folded and obtained pairing probabilities as described above, and finally calculated average pairing probability per position in this set and stored the result.

#### Structure context of Spb1 targets

For each 20-nt sequence surrounding either a target site or a background site, we used RNAsubopt (ViennaRNA package version 2.3.3) to obtain an ensemble of 100 suboptimal structures. These structures were labeled as follows: S for base-paired positions, B for unpaired positions within stems, L for positions in the loop of a hairpin loop, and F for unpaired positions at the ends of the sequence. For each position in the sequence, we calculated the frequency with which it assumes an S, F, L or B and used these as approximations of the probability that the position assumes each of these conformations.

#### GO Analyses

We used the statistical overrepresentation test in PANTHER (v13.0) to identify GO terms enriched in Spb1 target genes. As a background list of genes, we used all genes containing at least one site that met the MeTH-Seq read coverage threshold, regardless of peak height.

### DATA AND SOFTWARE AVAILABILITY

eCLIP-Seq and MeTH-Seq data may be downloaded from the Gene Expression Omnibus database, under accession number GSEXXXXXX. Analysis pipelines are available in <GitHub repository>.

## SUPPLEMENTAL FIGURE LEGENDS

**Figure S1.**
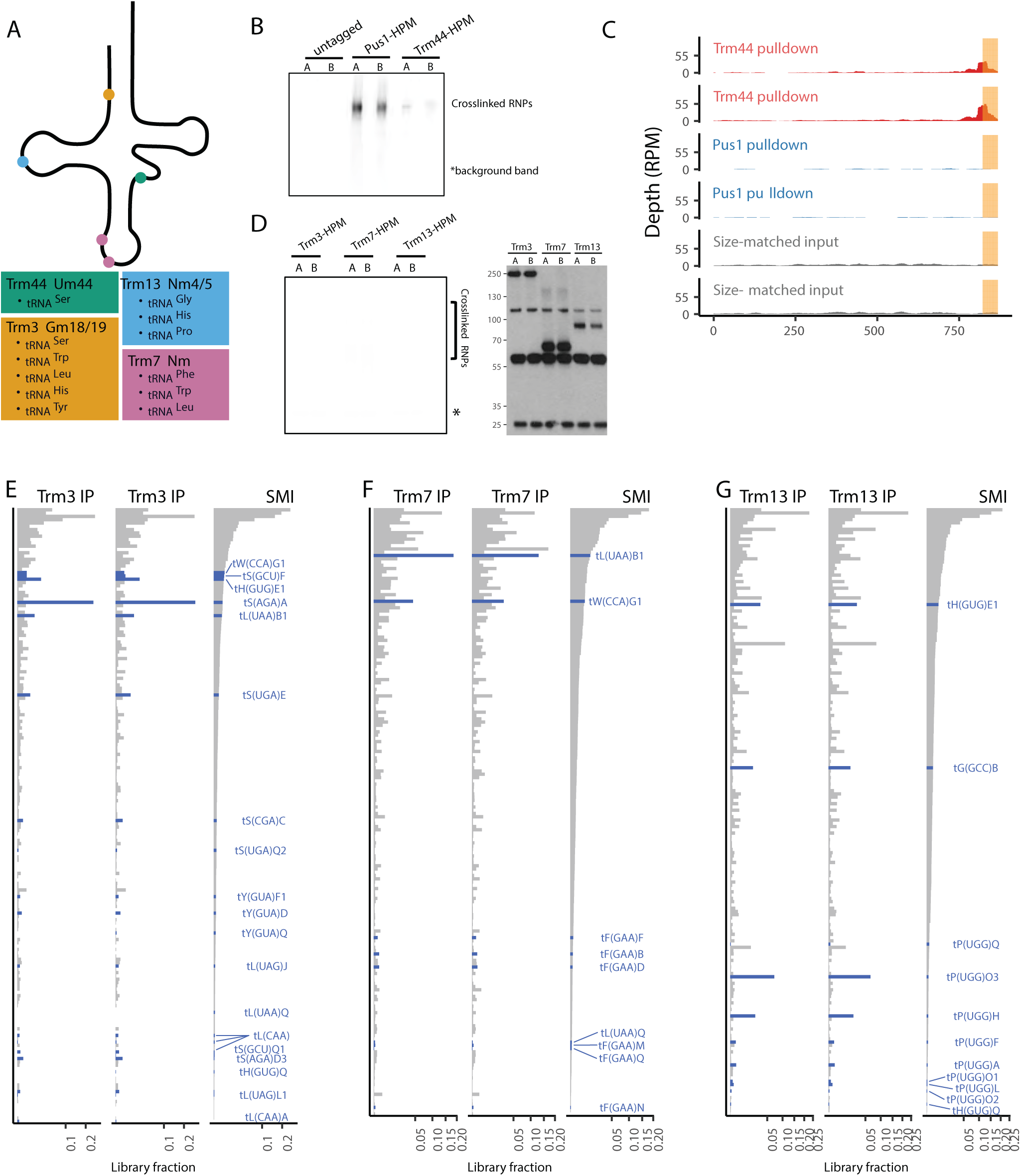
eCLIP Analysis of tRNA 2′-O-Methyltransferases, Related to Figure 1. (A)Summary of canonical non-coding RNA targets of yeast 2′-O-methyltransferases. (B)32P labeled RNA immunoprecipitated with anti-Myc from cross-linked Trm44-HPM and untagged control. (C)Read coverage in Pus1 and Trm44 eCLIP libraries for significantly enriched Trm44 mRNA eCLIP peaks (*p* ≤ 0.001 and ≥4-fold enriched versus SMI). (D)Anti-Myc immunoprecipitates from cross-linked Trm3-HPM, Trm7-HPM, and Trm13-HPM. Left: 32P labeled co-immunoprecipitated RNA. Right: Western blots. (E-G) Overlap between non-coding RNA eCLIP targets and canonical methylation substrates of Trm3 (E), Trm7 (F) and Trm13 (G). The fraction of the library mapped to each feature is shown for two eCLIP replicates and SMI control. Genes are ordered by their abundance in SMI. tRNAs that are known to be modified are highlighted in blue.

**Figure S2.**
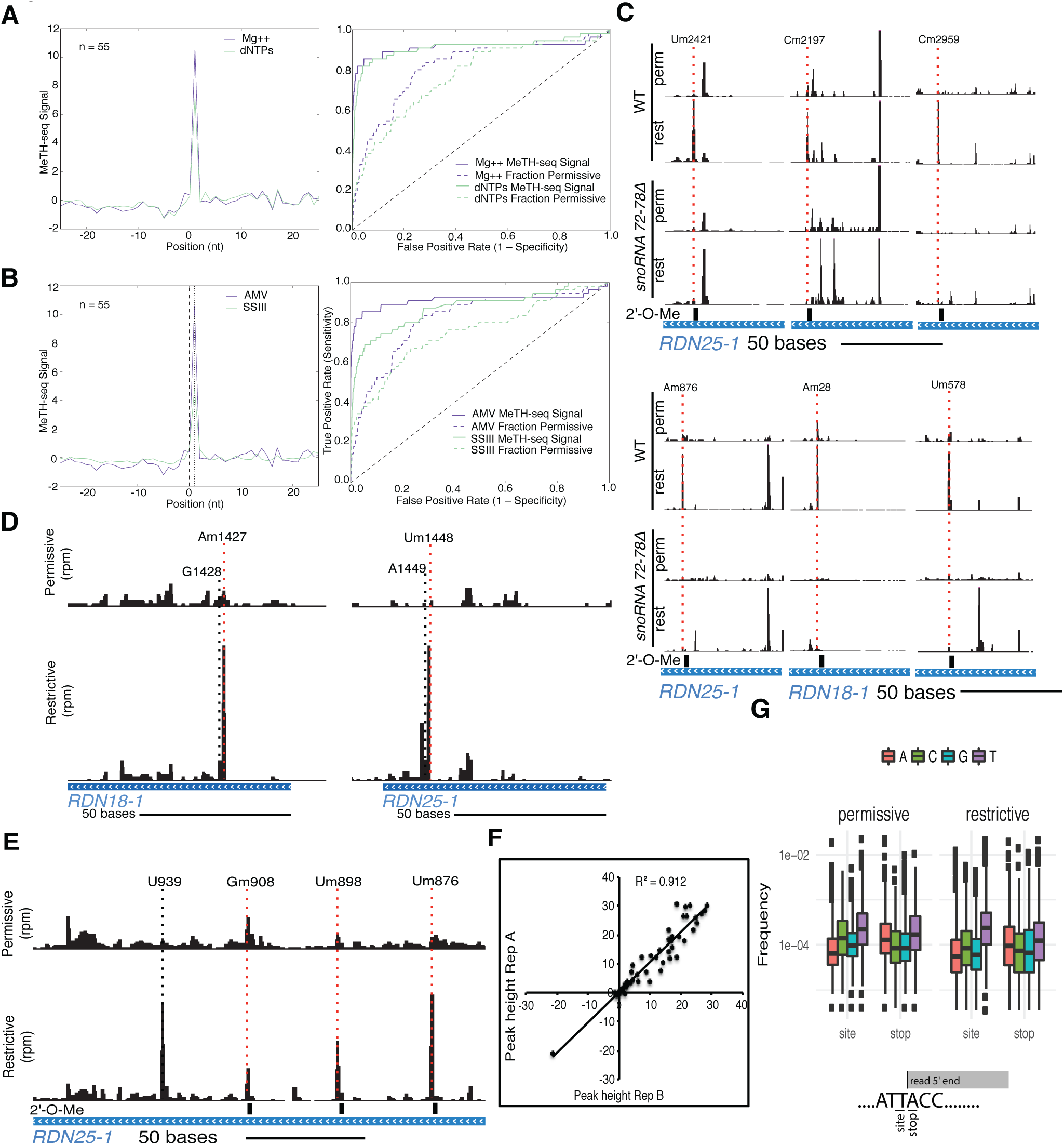
Controls for Transcriptome-wide Mapping of 2′-O-Methyl Ribose with MeTH-seq, Related to Figure 2. (A)Left: Meta plot of MeTH-seq signal at rRNA Nm sites detected by primer extension with limiting dNTPs (purple line) or limiting Mg^2^+ (green line). AMV was the reverse transcriptase used. Right: Receiver operating characteristic curves for MeTH-seq signal at all known Nm in rRNA obtained using different reaction conditions. (B)Left: Meta plot of MeTH-seq signal at rRNA Nm sites detected by primer extension using AMV (purple line) or SSIII (green line) reverse transcriptases. In both reactions,magnesium was limiting. Right: Receiver operating characteristic curves for MeTH-seq signal at all known Nm in rRNA obtained using different reverse transcriptases. (C)Deletion of C/D snoRNAs eliminated MeTH-seq peaks at their corresponding rRNA target sites. (D)Apparent Nm site misannotations identified by MeTH-seq. (E)Examples of false positive MeTH-seq peaks in rRNA. (F)Reproducibility of MeTH-seq peak heights at known Nm in rRNA. (G)Sequence bias in Nm site signal from MeTH-seq.

**Figure S3.**
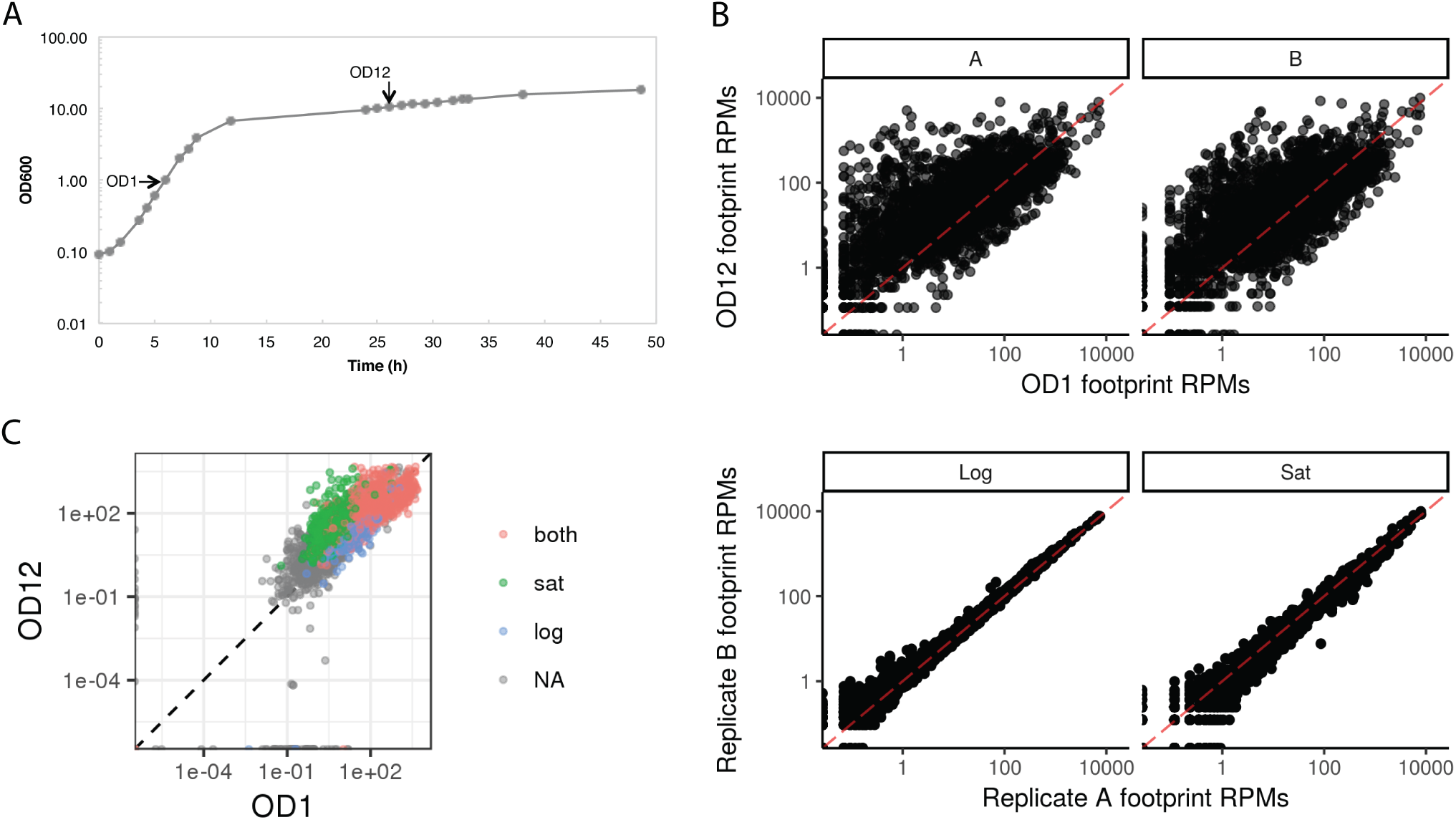
Conditions of Regulated mRNA 2′-O-Methylation, Related to Figure 3. (A)Yeast growth curve. Cells were harvested for MeTH-seq analysis in mid exponential growth (OD1) and when cultures reached OD12, after the diauxic shift and before stationary phase. (B)Ribosome footprint profiling reveals widespread differences in gene expression between log phase (OD1) and post-diauxic growth (OD12). Ribosome profiling results were reproducible, R^2^=0._-0._. Rpkm, reads per kilobase per million mapped reads. (C)RNA-seq data showing that expression levels minimally affect identification of regulated Nm sites in mRNAs. Average log-transformed expression levels (TPM) from two growth conditions for all mRNAs in which MeTH-seq identified Nm sites in both log phase and post-diauxic growth (red), log phase only (blue), and post-diauxic only (green). n=6 replicates for each condition.

**Figure S4.**
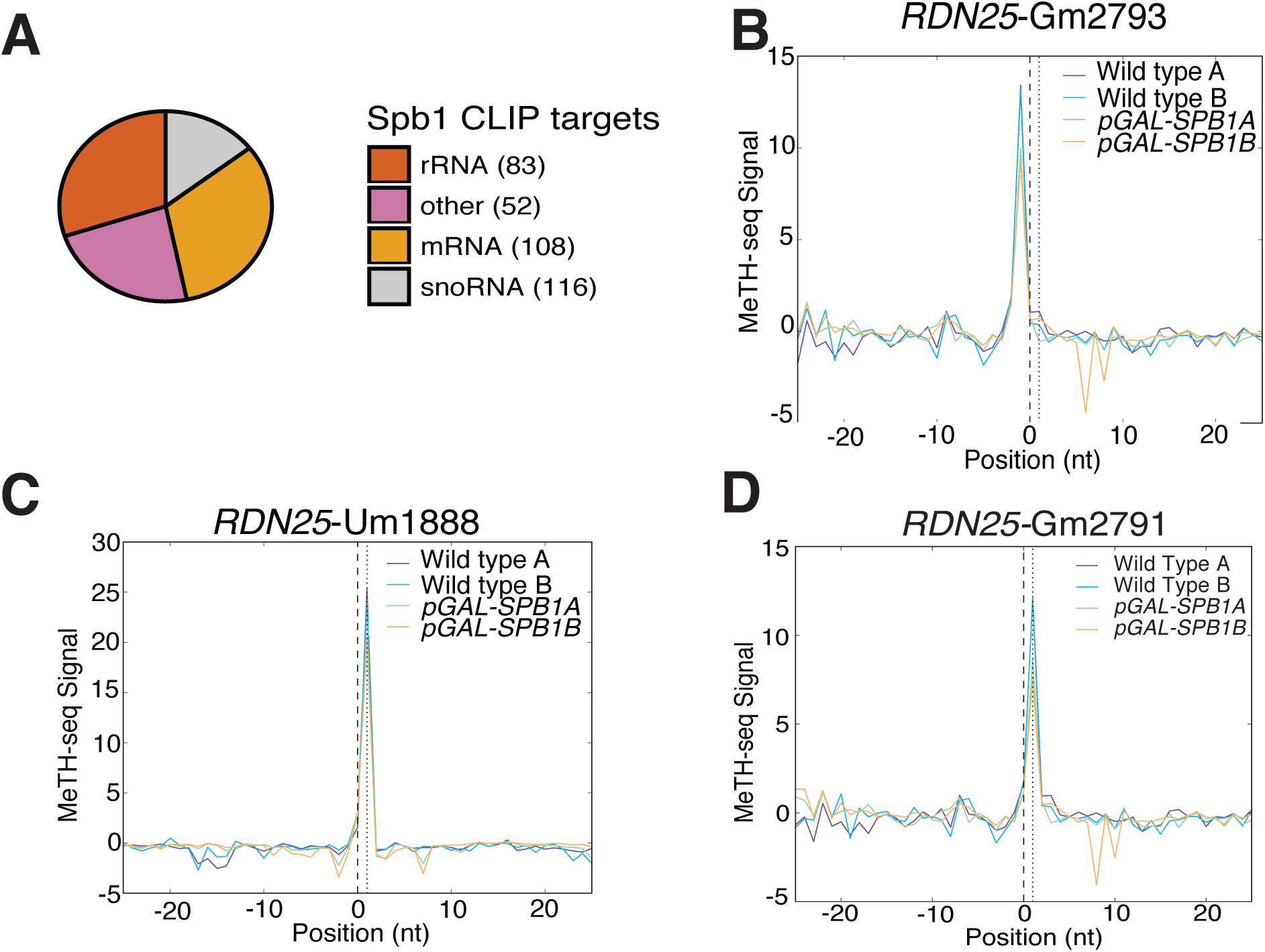
eCLIP Analysis of Spb1 and Effects of Spb1 Depletion on rRNA 2′-O-Methylation at snoRNA-Dependent Sites, Related to Figure 4. (A) Distribution of significantly enriched eCLIP peaks for Spb1, p ≤ 0.001 and ≥4-fold enriched versus SMI. (B-D) MeTH-seq signal plots of rRNA Nm sites with reproducible slight reductions of signal in *SPB1*-depleted cells. These sites are methylated by Nop1 guided by snR48 (B), snR62 (C), and snR48 (D).

**Figure S5.**
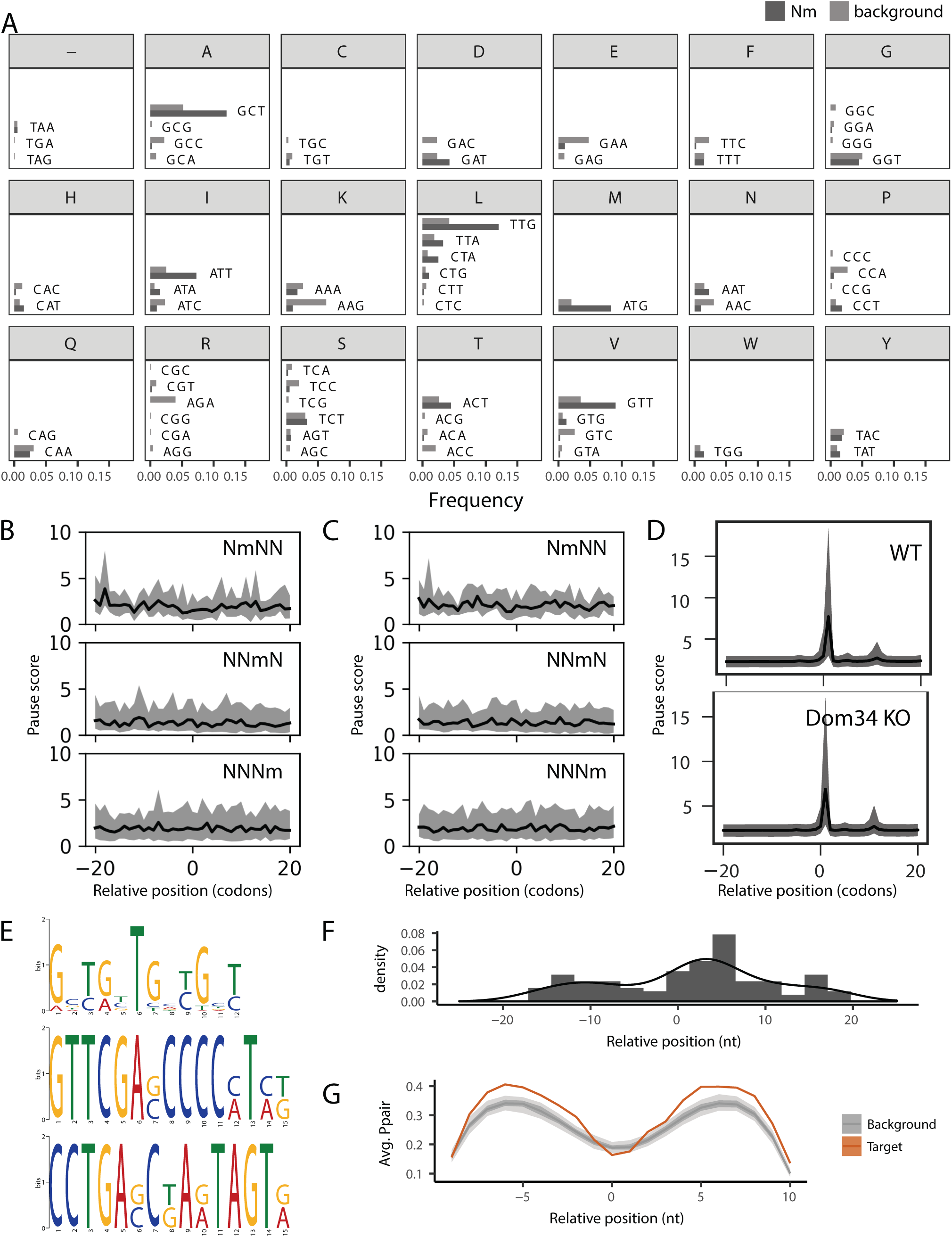
Codon and Motif Features of mRNA Target Sites of Spb1, Related to Figure 5. (A) Frequency of codons among *SPB1*-dependent Nm sites compared to the background of all codons with adequate read coverage for MeTH-seq analysis (Methods). (B-D) Ribosome footprint distributions. Ribosomal A sites aligned to Nm sites from wild type cells (B), *dom34Δ* mutants defective in No-Go Decay (C), and His codons in cells treated with 3-AT (D). Ribosome profiling data from (Guydosh and Green, 2014). (E)Web logos of significantly enriched motifs within a 50-nucleotide window surrounding novel *SPB1*-dependent Nm sites. The first motif (top) was found in 71/409 mRNA sites. The second was found exclusively in pre-tRNAs and the third was found in 15 mRNA sites. (F)Relative distances between *SPB1*-dependent Nm sites and the center of UGNUGN motifs (as called by MEME). (G)Average pairing probability in the 20-nt surrounding Spb1-dependent Nm sites (orange) or a background set of sites (grey).

**Figure S6.**
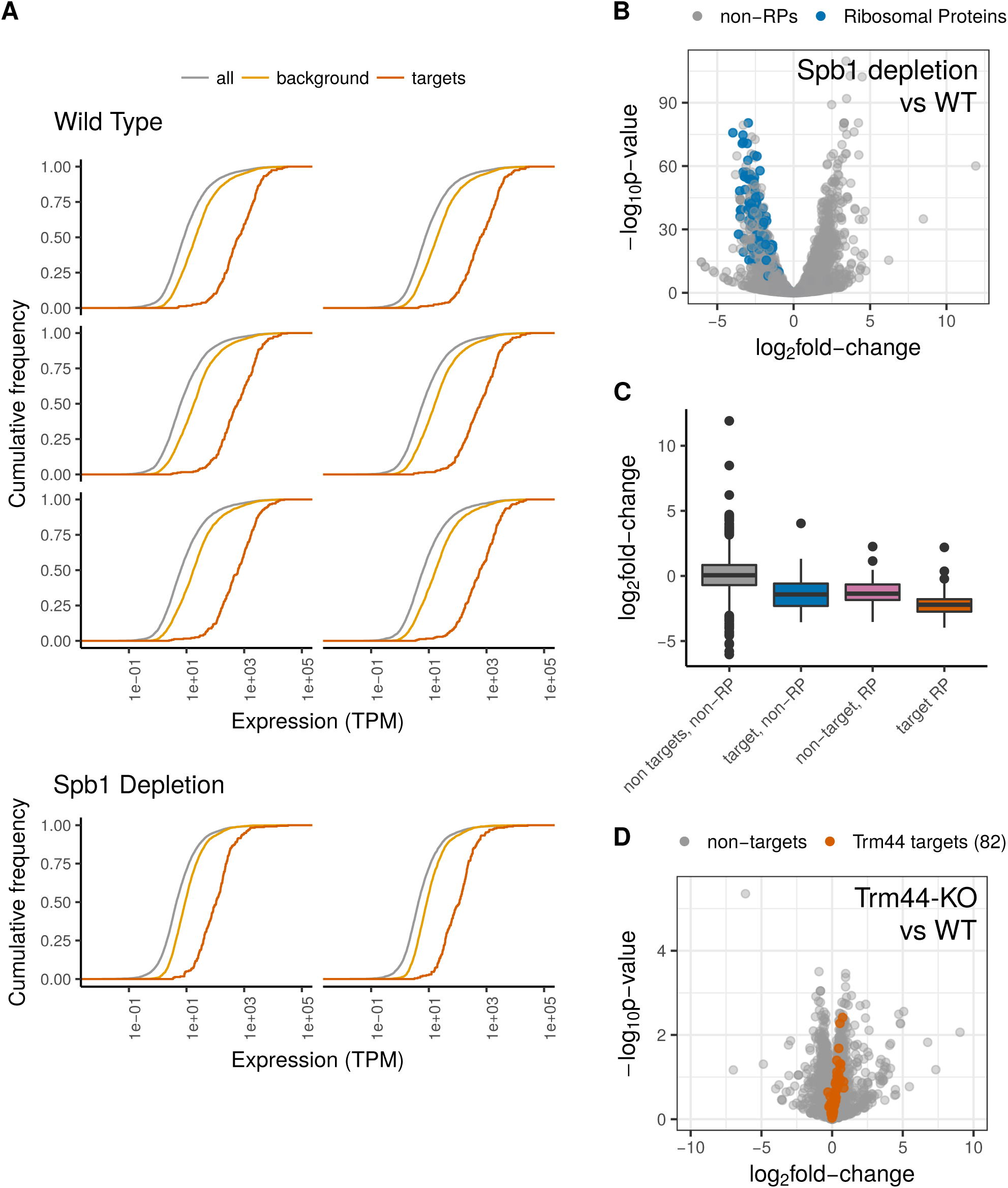
Global Effects of Spb1 Depletion on mRNA Abundance, Related to Figure 6. (A)Cumulative distributions of mRNA levels. Spb1 target genes were highly expressed compared to the background of all genes with adequate read coverage for MeTH-seq analysis in wild type controls (n=6) and showed reduced levels in Spb1-depleted cells (n=2). TPM, transcripts per million. KS test *p*-values are indicated. (B)Volcano plot of RNA-seq data shows reduced levels of ribosomal protein mRNAs in Spb1-depleted cells. (C)Levels of ribosomal protein mRNAs that are Spb1 targets were significantly reduced compared to non-targets. KS test, *p* < 5.9e-15.

## References

Bailey, T.L., Johnson, J., Grant, C.E., and Noble, W.S. (2015). The MEME Suite. Nucleic Acids Res. 43, W39–W49.

Baltz, A.G., Munschauer, M., Schwanhäusser, B., Vasile, A., Murakawa, Y., Schueler, M., Youngs, N., Penfold-Brown, D., Drew, K., Milek, M., et al. (2012). The mRNA-Bound Proteome and Its Global Occupancy Profile on Protein-Coding Transcripts. Mol. Cell 46, 674–690.

Beckmann, B.M. (2017). RNA interactome capture in yeast. Methods 118-119, 82–92.

Beckmann, B.M., Horos, R., Fischer, B., Castello, A., Eichelbaum, K., Alleaume, A.-M., Schwarzl, T., Curk, T.T., Foehr, S., Huber, W., et al. (2015). The RNA-binding proteomes from yeast to man harbour conserved enigmRBPs. Nat. Commun. 6, 10127.

Bonnerot, C., Pintard, L., and Lutfalla, G. (2003). Functional redundancy of Spb1p and a snR52-dependent mechanism for the 2′-O-ribose methylation of a conserved rRNA position in yeast. Mol. Cell 12, 1309–1315.

Brar, G. a., Yassour, M., Friedman, N., Regev, A., Ingolia, N.T., and Weissman, J.S. (2012). High-Resolution View of the Yeast Meiotic Program Revealed by Ribosome Profiling. Science (80-.). 335, 552–557.

Breker, M., Gymrek, M., and Schuldiner, M. (2013). A novel single-cell screening platform reveals proteome plasticity during yeast stress responses. J. Cell Biol. 200, 839–850.

Butcher, S.E., and Pyle, A.M. (2011). The molecular interactions that stabilize RNA tertiary structure: RNA motifs, patterns, and networks. Acc. Chem. Res. 44, 1302–1311.

Carlile, T.M., Rojas-Duran, M.F., Zinshteyn, B., Shin, H., Bartoli, K.M., and Gilbert, W. V (2014). Pseudouridine profiling reveals regulated mRNA pseudouridylation in yeast and human cells. Nature 515, 143–146.

Carlile, T.M., Rojas-Duran, M.F., and Gilbert, W. V. (2015). Transcriptome-Wide Identification of Pseudouridine Modifications Using Pseudo-seq. Curr. Protoc. Mol. Biol. 112, 4.25.1–4.25.24.

Castello, A., Fischer, B., Eichelbaum, K., Horos, R., Beckmann, B., Strein, C., Davey, N., Humphreys, D., Preiss, T., Steinmetz, L., et al. (2012). Insights into RNA Biology from an Atlas of Mammalian mRNA-Binding Proteins. Cell 149, 1393–1406.

Cavaillé, J., Chetouani, F., and Bachellerie, J.P. (1999). The yeast Saccharomyces cerevisiae YDL112w ORF encodes the putative 2’-O-ribose methyltransferase catalyzing the formation of Gm18 in tRNAs. RNA 5, 66–81.

Conway, A.E., Van Nostrand, E.L., Pratt, G.A., Aigner, S., Wilbert, M.L., Sundararaman, B., Freese, P., Lambert, N.J., Sathe, S., Liang, T.Y., et al. (2016). Enhanced CLIP Uncovers IMP Protein-RNA Targets in Human Pluripotent Stem Cells Important for Cell Adhesion and Survival. Cell Rep. 15, 666–679.

Coyle, S.M., Gilbert, W. V., Doudna, J.A., Coyle SM, Gilbert WV, and Doudna JA (2009). Direct link between RACK1 function and localization at the ribosome in vivo. Mol. Cell. Biol. 29, 1626–1634.

Czauderna, F., Fechtner, M., Dames, S., Aygün, H., Klippel, A., Pronk, G.J., Giese, K., and Kaufmann, J. (2003). Structural variations and stabilising modifications of synthetic siRNAs in mammalian cells. Nucleic Acids Res. 31, 2705–2716.

Dai, Q., Moshitch-Moshkovitz, S., Han, D., Kol, N., Amariglio, N., Rechavi, G., Dominissini, D., and He, C. (2017). Nm-seq maps 2′-O-methylation sites in human mRNA with base precision. Nat. Methods 14, 695–698.

Devarkar, S.C., Wang, C., Miller, M.T., Ramanathan, A., Jiang, F., Khan, A.G., Patel, S.S., and Marcotrigiano, J. (2016). Structural basis for m7G recognition and 2′-O-methyl discrimination in capped RNAs by the innate immune receptor RIG-I. Proc. Natl. Acad. Sci. 113, 596–601.

Dunlap, B.E., Friderici, K.H., and Rottman, F. (1971). 2’-O-Methyl polynucleotides as templates for cell-free amino acid incorporation. Biochemistry 10, 2581–2587.

Ghaemmaghami, S., Huh, W.-K., Bower, K., Howson, R., Bower, K., Belle, A., Howson, R.W., Belle, A., Dephoure, N., O’Shea, E., et al. (2003). Global analysis of protein expression in yeast. Nature 425, 737–741.

Gilbert, W. V., Bell, T.A., and Schaening, C. (2016). Messenger RNA modifications: Form, distribution, and function. Science (80-.). 352, 1408–1412.

Gillen, A.E., Yamamoto, T.M., Kline, E., Hesselberth, J.R., and Kabos, P. (2016). Improvements to the HITS-CLIP protocol eliminate widespread mispriming artifacts. BMC Genomics 17, 338.

Gonzalo, P., and Reboud, J.P. (2003). The puzzling lateral flexible stalk of the ribosome. Biol. Cell 95, 179–193.

Graumann, J., Dunipace, L.A., Seol, J.H., McDonald, W.H., Yates, J.R., Wold, B.J., and Deshaies, R.J. (2004). Applicability of tandem affinity purification MudPIT to pathway proteomics in yeast. Mol. Cell. Proteomics 3, 226–237.

Guy, M.P., and Phizicky, E.M. (2014). Two-subunit enzymes involved in eukaryotic post-transcriptional tRNA modification. RNA Biol. 11, 1608–1618.

Guydosh, N.R., and Green, R. (2014). Dom34 Rescues Ribosomes in 3′ Untranslated Regions. Cell 156, 950–962.

Hanson, G., and Coller, J. (2017). Codon optimality, bias and usage in translation and mRNA decay. Nat. Rev. Mol. Cell Biol.

Harnpicharnchai, P., Jakovljevic, J., Horsey, E., Miles, T., Roman, J., Rout, M., Meagher, D., Imai, B., Guo, Y., Brame, C.J., et al. (2001). Composition and functional characterization of yeast 66S ribosome assembly intermediates. Mol. Cell 8, 505–515.

Hoernes, T.P., Clementi, N., Faserl, K., Glasner, H., Breuker, K., Lindner, H., Hüttenhofer, A., and Erlacher, M.D. (2015). Nucleotide modifications within bacterial messenger RNAs regulate their translation and are able to rewire the genetic code. Nucleic Acids Res.

Hou, Y.M., Zhang, X., Holland, J.A., and Davis, D.R. (2001). An important 2’-OH group for an RNA-protein interaction. Nucleic Acids Res. 29, 976–985.

Incarnato, D., Anselmi, F., Morandi, E., Neri, F., Maldotti, M., Rapelli, S., Parlato, C., Basile, G., and Oliviero, S. (2017). High-throughput single-base resolution mapping of RNA 2?-O-methylated residues. Nucleic Acids Res.

Inoue, H., Hayase, Y., Imura, A., Iwai, S., Miura, K., and Ohtsuka, E. (1987). Synthesis and hybridization studies on two complementary nona(2’-o-methyl)ribonucleotides. Nucleic Acids Res. 15, 6131–6148.

Jones, S., Daley, D.T., Luscombe, N.M., Berman, H.M., and Thornton, J.M. (2001). Protein-RNA interactions: a structural analysis. Nucleic Acids Res. 29, 943–954.

Kawai, G., Yamamoto, Y., Kamimura, T., Masegi, T., Sekine, M., Hata, T., Iimori, T., Watanabe, T., Miyazawa, T., and Yokoyama, S. (1992). Conformational rigidity of specific pyrimidine residues in tRNA arises from posttranscriptional modifications that enhance steric interaction between the base and the 2’-hydroxyl group. Biochemistry 31, 1040–1046.

Ke, S., Pandya-Jones, A., Saito, Y., Fak, J.J., Vågbø, C.B., Geula, S., Hanna, J.H., Black, D.L., Darnell, J.E., and Darnell, R.B. (2017). m ^6^ A mRNA modifications are deposited in nascent pre-mRNA and are not required for splicing but do specify cytoplasmic turnover. Genes Dev. 31, 990–1006.

Keffer-Wilkes, L.C., Veerareddygari, G.R., and Kothe, U. (2016). RNA modification enzyme TruB is a tRNA chaperone. Proc. Natl. Acad. Sci. 113, 14306–14311.

Kiss, T. (2002). Small nucleolar RNAs: an abundant group of noncoding RNAs with diverse cellular functions. Cell 109, 145–148.

Kiss-László, Z., Henry, Y., Bachellerie, J.P., Caizergues-Ferrer, M., and Kiss, T. (1996). Site-specific ribose methylation of preribosomal RNA: a novel function for small nucleolar RNAs. Cell 85, 1077–1088.

Kotelawala, L., Grayhack, E.J., and Phizicky, E.M. (2007). Identification of yeast tRNA Um44 2’-O-methyltransferase (Trm44) and demonstration of a Trm44 role in sustaining levels of specific tRNASer species. RNA 14, 158–169.

Kotelawala, L., Grayhack, E.J., and Phizicky, E.M. (2008). Identification of yeast tRNA Um(44) 2’-O-methyltransferase (Trm44) and demonstration of a Trm44 role in sustaining levels of specific tRNA(Ser) species. RNA 14, 158–169.

Kressler, D., Rojo, M., Linder, P., and Cruz, J. (1999). Spb1p is a putative methyltransferase required for 60S ribosomal subunit biogenesis in Saccharomyces cerevisiae. Nucleic Acids Res. 27, 4598–4608.

Krogh, N., Jansson, M.D., H?fner, S.J., Tehler, D., Birkedal, U., Christensen-Dalsgaard, M., Lund, A.H., Nielsen, H., Häfner, S.J., Tehler, D., et al. (2016). Profiling of 2’-O-Me in human rRNA reveals a subset of fractionally modified positions and provides evidence for ribosome heterogeneity. Nucleic Acids Res. 44.

Kulak, N.A., Pichler, G., Paron, I., Nagaraj, N., and Mann, M. (2014). Minimal, encapsulated proteomic-sample processing applied to copy-number estimation in eukaryotic cells. Nat. Methods 11, 319–324.

Lacoux, C., Di Marino, D., Boyl, P.P., Zalfa, F., Yan, B., Ciotti, M.T., Falconi, M., Urlaub, H., Achsel, T., Mougin, A., et al. (2012). BC1-FMRP interaction is modulated by 2′-O-methylation: RNA-binding activity of the tudor domain and translational regulation at synapses. Nucleic Acids Res. 40, 4086–4096.

Lamm, A.T., Stadler, M.R., Zhang, H., Gent, J.I., and Fire, A.Z. (2011). Multimodal RNA-seq using single-strand, double-strand, and CircLigase-based capture yields a refined and extended description of the C. elegans transcriptome. Genome Res. 21, 265–275.

Lapeyre, B., and Purushothaman, S.K. (2004). Spb1p-directed formation of Gm2922 in the ribosome catalytic center occurs at a late processing stage. Mol. Cell 16, 663–669.

Lavoie, M., and Abou Elela, S. (2008). Yeast Ribonuclease III Uses a Network of Multiple Hydrogen Bonds for RNA Binding and Cleavage ^†^. Biochemistry 47, 8514–8526.

Lebars, I., Legrand, P., Aimé, A., Pinaud, N., Fribourg, S., and Di Primo, C. (2008). Exploring TAR-RNA aptamer loop-loop interaction by X-ray crystallography, UV spectroscopy and surface plasmon resonance. Nucleic Acids Res. 36, 7146–7156.

Li, X., Xiong, X., and Yi, C. (2016). Epitranscriptome sequencing technologies: decoding RNA modifications. Nat. Methods 14, 23–31.

Longtine, M.S., McKenzie, A., Demarini, D.J., Shah, N.G., Wach, A., Brachat, A., Philippsen, P., Pringle, J.R., Longtine MS, McKenzie A, et al. (1998). Additional modules for versatile and economical PCR-based gene deletion and modification in Saccharomyces cerevisiae. Yeast 14, 953–961.

Lovci, M.T., Ghanem, D., Marr, H., Arnold, J., Gee, S., Parra, M., Liang, T.Y., Stark, T.J., Gehman, L.T., Hoon, S., et al. (2013). Rbfox proteins regulate alternative mRNA splicing through evolutionarily conserved RNA bridges. Nat. Struct. Mol. Biol. 20, 1434–1442.

Lovejoy, A.F., Riordan, D.P., and Brown, P.O. (2014). Transcriptome-wide mapping of pseudouridines: pseudouridine synthases modify specific mRNAs in S. cerevisiae. PLoS One 9, e110799.

Lowe, T.M., and Eddy, S.R. (1997). tRNAscan-SE: a program for improved detection of transfer RNA genes in genomic sequence. Nucleic Acids Res. 25, 955–964.

Machnicka, M.A., Milanowska, K., Osman Oglou, O., Purta, E., Kurkowska, M., Olchowik, A., Januszewski, W., Kalinowski, S., Dunin-Horkawicz, S., Rother, K.M., et al. (2013). MODOMICS: a database of RNA modification pathways--2013 update. Nucleic Acids Res. 41, D262–7.

Maden, B.E., Corbett, M.E., Heeney, P.A., Pugh, K., and Ajuh, P.M. (1995). Classical and novel approaches to the detection and localization of the numerous modified nucleotides in eukaryotic ribosomal RNA. Biochimie 77, 22–29.

Majlessi, M., Nelson, N.C., and Becker, M.M. (1998). Advantages of 2’-O-methyl oligoribonucleotide probes for detecting RNA targets. Nucleic Acids Res. 26, 2224–2229.

Marchand, V., Blanloeil-Oillo, F., Helm, M., and Motorin, Y. (2016). Illumina-based RiboMethSeq approach for mapping of 2′-O-Me residues in RNA. Nucleic Acids Res. 44, e135–e135.

Morello, L.G., Coltri, P.P., Quaresma, A.J.C., Simabuco, F.M., Silva, T.C.L., Singh, G., Nickerson, J.A., Oliveira, C.C., Moore, M.J., and Zanchin, N.I.T. (2011). The Human Nucleolar Protein FTSJ3 Associates with NIP7 and Functions in Pre-rRNA Processing. PLoS One 6, e29174.

Motorin, Y., Muller, S., Behm-Ansmant, I., and Branlant, C. (2007). Identification of Modified Residues in RNAs by Reverse Transcription-Based Methods. In Methods in Enzymology, pp. 21–53.

Müller, S., Windhof, I.M., Maximov, V., Jurkowski, T., Jeltsch, A., Förstner, K.U., Sharma, C.M., Gräf, R., and Nellen, W. (2013). Target recognition, RNA methylation activity and transcriptional regulation of the Dictyostelium discoideum Dnmt2-homologue (DnmA). Nucleic Acids Res. 41, 8615–8627.

Nicoloso, M., Qu, L.-H., Michot, B., and Bachellerie, J.-P. (1996). Intron-encoded, Antisense Small Nucleolar RNAs: The Characterization of Nine Novel Species Points to Their Direct Role as Guides for the 2′-O-ribose Methylation of rRNAs. J. Mol. Biol. 260, 178–195.

Van Nostrand, E.L., Pratt, G.A., Shishkin, A.A., Gelboin-Burkhart, C., Fang, M.Y., Sundararaman, B., Blue, S.M., Nguyen, T.B., Surka, C., Elkins, K., et al. (2016). Robust transcriptome-wide discovery of RNA-binding protein binding sites with enhanced CLIP (eCLIP). Nat. Methods 13, 1–9.

Pelechano, V., Wei, W., Jakob, P., and Steinmetz, L.M. (2014). Genome-wide identification of transcript start and end sites by transcript isoform sequencing. Nat. Protoc. 9, 1740–1759.

Phizicky, E.M., and Hopper, A.K. (2010). tRNA biology charges to the front. Genes Dev. 24, 1832–1860.

Piekna-Przybylska, D., Decatur, W.A., and Fournier, M.J. (2007). New bioinformatic tools for analysis of nucleotide modifications in eukaryotic rRNA. RNA 13, 305–312.

Pintard, L., Lecointe, F., Bujnicki, J.M., Bonnerot, C., Grosjean, H., and Lapeyre, B. (2002a). Trm7p catalyses the formation of two 2’-O-methylriboses in yeast tRNA anticodon loop. EMBO J. 21, 1811–1820.

Pintard, L., Bujnicki, J.M., Lapeyre, B., and Bonnerot, C. (2002b). MRM2 encodes a novel yeast mitochondrial 21s rRNA methyltransferase. EMBO J. 21, 1139–1147.

Raabe, C.A., Tang, T.-H., Brosius, J., and Rozhdestvensky, T.S. (2014). Biases in small RNA deep sequencing data. Nucleic Acids Res. 42, 1414–1426.

Roignant, J.-Y., and Soller, M. (2017). m 6 A in mRNA: An Ancient Mechanism for Fine-Tuning Gene Expression. Trends Genet. 33, 380–390.

Sachs, A.B., Davis, R.W., and Kornberg, R.D. (1987). A Single Domain of Yeast Poly (A)-Binding Protein Is Necessary and Sufficient for RNA Binding and Cell Viability. 7, 3268–3276.

Schaefer, M., Kapoor, U., and Jantsch, M.F. (2017). Understanding RNA modifications: the promises and technological bottlenecks of the ?epitranscriptome? Open Biol. 7, 170077.

Schwartz, S., Agarwala, S.D., Mumbach, M.R., Jovanovic, M., Mertins, P., Shishkin, A., Tabach, Y., Mikkelsen, T.S., Satija, R., Ruvkun, G., et al. (2013). High-resolution mapping reveals a conserved, widespread, dynamic mRNA methylation program in yeast meiosis. Cell 155, 1409–1421.

Schwartz, S., Bernstein, D.A., Mumbach, M.R., Jovanovic, M., Herbst, R.H., León-Ricardo, B.X., Engreitz, J.M., Guttman, M., Satija, R., Lander, E.S., et al. (2014). Transcriptome-wide Mapping Reveals Widespread Dynamic-Regulated Pseudouridylation of ncRNA and mRNA. Cell 159, 148–162.

Simabuco, F.M., Morello, L.G., Aragão, A.Z.B., Paes Leme, A.F., and Zanchin, N.I.T. (2012). Proteomic Characterization of the Human FTSJ3 Preribosomal Complexes. 11.

Simon, B., Kirkpatrick, J.P., Eckhardt, S., Reuter, M., Rocha, E.A., Andrade-Navarro, M.A., Sehr, P., Pillai, R.S., and Carlomagno, T. (2011). Recognition of 2′-O-Methylated 3′-End of piRNA by the PAZ Domain of a Piwi Protein. Structure 19, 172–180.

Sirum-Connolly, K., and Mason, T.L. (1993). Functional requirement of a site-specific ribose methylation in ribosomal RNA. Science 262, 1886–1889.

Sproat, B.S., Lamond, A.I., Beijer, B., Neuner, P., and Ryder, U. (1989). Highly efficient chemical synthesis of 2???-O-methyloligoribonucleotides and tetrabiotinylated derivatives; novel probes that are resistant to degradation by RNA or DNA specific nucleases. Nucleic Acids Res. 17, 3373–3386.

Stanley, S.E., Gable, D.L., Wagner, C.L., Carlile, T.M., Hanumanthu, V.S., Podlevsky, J.D., Khalil, S.E., DeZern, A.E., Rojas-Duran, M.F., Applegate, C.D., et al. (2016). Loss-of-function mutations in the RNA biogenesis factor NAF1 predispose to pulmonary fibrosis-emphysema. Sci. Transl. Med. 8, 351ra107.

Tian, Y., Simanshu, D.K., Ma, J.-B., and Patel, D.J. (2011). Structural basis for piRNA 2’-O-methylated 3’-end recognition by Piwi PAZ (Piwi/Argonaute/Zwille) domains. Proc. Natl. Acad. Sci. 108, 903–910.

Treger, M., and Westhof, E. (2001). Statistical analysis of atomic contacts at RNA-protein interfaces. J. Mol. Recognit. 14, 199–214.

Tsourkas, A., Behlke, M.A., and Bao, G. (2002). Hybridization of 2′-O-methyl and 2-deoxy molecular beacons to RNA and DNA targets. Nucleic Acids Res. 30, 5168–5174.

Wang, X., Lu, Z., Gomez, A., Hon, G.C., Yue, Y., Han, D., Fu, Y., Parisien, M., Dai, Q., Jia, G., et al. (2014). N6-methyladenosine-dependent regulation of messenger RNA stability. Nature 505, 117–120.

Wilkinson, M.L., Crary, S.M., Jackman, J.E., Grayhack, E.J., and Phizicky, E.M. (2007). The 2’-O-methyltransferase responsible for modification of yeast tRNA at position 4. RNA 13, 404–413.

Winzeler, E.A., Shoemaker, D.D., Astromoff, A., Liang, H., Anderson, K., Andre, B., Bangham, R., Benito, R., Boeke, J.D., Bussey, H., et al. (1999). Functional characterization of the S. cerevisiae genome by gene deletion and parallel analysis. Science 285, 901–906.

Wlodarski, T., Kutner, J., Towpik, J., Knizewski, L., Rychlewski, L., Kudlicki, A., Rowicka, M., Dziembowski, A., and Ginalski, K. (2011). Comprehensive Structural and Substrate Specificity Classification of the Saccharomyces cerevisiae Methyltransferome. PLoS One 6, e23168.

Yang, J., Sharma, S., Kötter, P., and Entian, K.-D. (2015). Identification of a new ribose methylation in the 18S rRNA of S. cerevisiae. Nucleic Acids Res. 43, 2342–2352.

Yue, Y., Liu, J., and He, C. (2015). RNA N 6-methyladenosine methylation in post-transcriptional gene expression regulation. Genes Dev. 29, 1343–1355.

